# Discrete Correlate Summation Clustering of a Dopamine-Angiotensin Network

**DOI:** 10.1101/2024.06.22.600163

**Authors:** Brian M. Westwood, Liliya M. Yamaleyeva, Rong Chen

## Abstract

We present a model of directed investigation of KEGG pathway analysis, beginning with discrete correlate summation (DCS) clustering of Swiss-Prot sets. Dopamine and angiotensin related proteins define the boundary for pathway analyses upon a network of proteins also associated with a compositely defined matrix of Swiss-Prot keywords to provide a framework for their biological indices (Figure 1).

## INTRODUCTION

This study is a framework. It is designed to visualize a system between two disparate topics, connected by common relationships.

Generally, we categorize genes with differential expression between treatments, to investigate enrichment of an effected gene set, and decide if there is evidence to indicate a concerted alteration of a pathway and not just stochastic changes.

This framework is constructed using the classification structure of the Kyoto Encyclopedia of Genes and Genomes (KEGG) (Kanehisa et al., 2000) molecular network database, evaluated with the ConsensusPathDB-human (CPDB) (Kamburov et al., 2011) over-representation analysis, using very specific starting sets (Swiss-Prot keyword searches) (UniProt Consortium, 2021).

## METHODS

This is an investigation of 3309 proteins from the 20 SwissProt keyword searches [oxidative&stress, glutamate, inflammation, memory, glutamatergic, blood&pressure, dopamine, hypertension, blood&brain&barrier, gaba, angiotensin, prefrontal&cortex, dopaminergic, gabaergic, cognition, midbrain, cerebral&vascular, striatum, renin, ageing].

Iterative ‘original matrix total array average was calculated right to left’ (RL-OriMaTArA) DCS clustering (Bronson et al., 2022) yields 3 clear count domains and 2 singleton searches for the 3309 proteins (Figure 2a, 2b). Briefly, upper cluster of a matrix was determined by sorting the totals of each variable column, then repeating on remaining extra-cluster submatrix.

**Figure 1.**
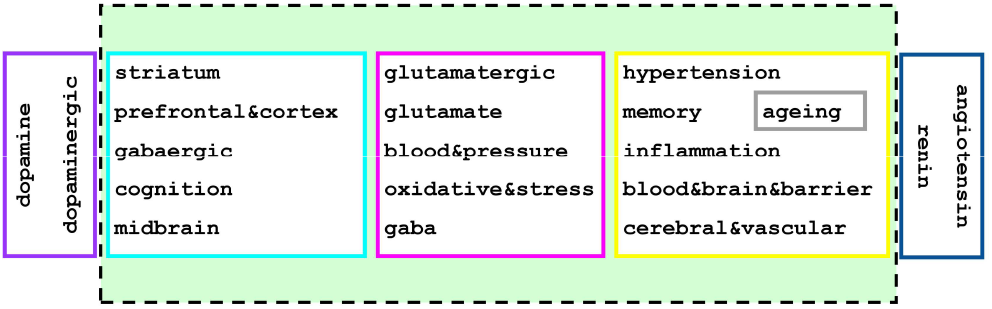
Graphical abstract

**Figure 2.**
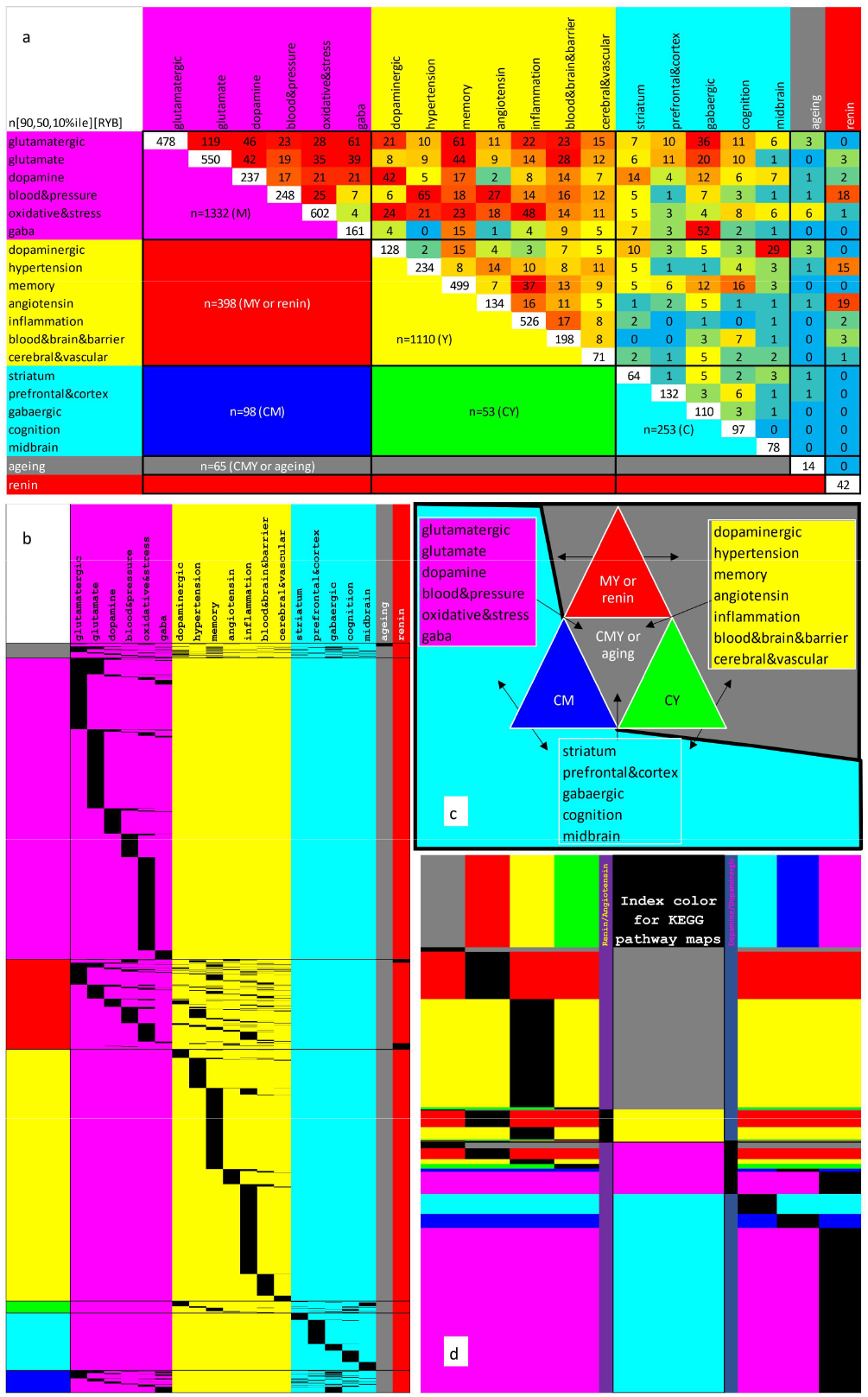
Binary count clustering for Swiss-Prot keyword searches (a); keyword group membership columns by cluster row groups [(b) rows are individual proteins, black cell = membership]; ternary diagram of cluster layout [(c) with underlay of node coloring for associated groups]; groups of clusters organized by index color for KEGG pathway map contingency tests [(d) RAS associated genes (gray) RAS genes (yellow) DA genes (magenta) DA associated genes (cyan)].

RL-OriMaTArA produced 3 primary clusters [C, M, Y] and 4 combination sets [CM-blue, MY-red, CY-green, CMY-gray] to search in CPDB for pathway enrichment analysis (Figure 2c, 2d) of KEGG pathways.

The 7 CPDB searches of 3309 proteins yielded a primary output of 137 KEGG pathways with 1991 genes. Area under the curve (AUC) of receiver operating characteristic (ROC) curves and precision-recall F-measures (Fawcett, 2006) were calculated for neighborhoods 1-5 for 51 KEGG pathways and 1045 genes with 4 or more angiotensin or renin Swiss-Prot search proteins/genes and with 4 or more dopamine or dopaminergic Swiss-Prot search proteins/genes in each pathway (Table 1).

**Table 1.** KEGG pathway contingency test output, sorted by TP then maximum for AUC of ROC curve and F-measure.

| KEGG | Pathway | AUC | F | TP |
| --- | --- | --- | --- | --- |
| hsa04270 | Vascular smooth muscle contraction |  | 0.96 | 12 |
| hsa04925 | Aldosterone synthesis and secretion |  | 1.00 | 10 |
| hsa04022 | cGMP-PKG signaling pathway | 0.80 | 0.90 | 9 |
| hsa04071 | Sphingolipid signaling pathway |  | 1.00 | 8 |
| hsa04810 | Regulation of actin cytoskeleton |  | 1.00 | 8 |
| hsa04927 | Cortisol synthesis and secretion |  | 1.00 | 8 |
| hsa04750 | Inflammatory mediator regulation of TRP channels |  | 0.94 | 8 |
| hsa04611 | Platelet activation |  | 1.00 | 7 |
| hsa04024 | cAMP signaling pathway | 0.96 | 0.93 | 7 |
| hsa04510 | Focal adhesion |  | 0.93 | 7 |
| hsa04921 | Oxytocin signaling pathway | 0.89 | 0.88 | 7 |
| hsa04062 | Chemokine signaling pathway | 0.75 | 0.88 | 7 |
| hsa04020 | Calcium signaling pathway | 1.00 | 1.00 | 6 |
| hsa04066 | HIF-1 signaling pathway |  | 1.00 | 6 |
| hsa04151 | PI3K-Akt signaling pathway |  | 1.00 | 6 |
| hsa04360 | Axon guidance | 0.99 | 0.92 | 6 |
| hsa04072 | Phospholipase D signaling pathway | 0.97 | 0.92 | 6 |
| hsa04015 | Rap1 signaling pathway | 0.75 | 0.92 | 6 |
| hsa04924 | Renin secretion |  | 0.92 | 6 |
| hsa04926 | Relaxin signaling pathway | 0.86 | 0.92 | 6 |
| hsa04261 | Adrenergic signaling in cardiomyocytes |  | 1.00 | 5 |
| hsa04728 | Dopaminergic synapse |  | 1.00 | 5 |
| hsa04911 | Insulin secretion |  | 1.00 | 5 |
| hsa04913 | Ovarian steroidogenesis |  | 1.00 | 5 |
| hsa04010 | MAPK signaling pathway | 0.87 | 0.83 | 5 |
| hsa04670 | Leukocyte transendothelial migration | 0.40 | 0.83 | 5 |
| hsa04720 | Long-term potentiation |  | 1.00 | 4 |
| hsa04725 | Cholinergic synapse |  | 1.00 | 4 |
| hsa04310 | Wnt signaling pathway | 0.81 | 0.89 | 4 |
| hsa04726 | Serotonergic synapse | 0.75 | 0.89 | 4 |
| hsa04928 | Parathyroid hormone synthesis, secretion and action | 0.81 | 0.89 | 4 |
| hsa04912 | GnRH signaling pathway | 0.00 | 0.80 | 4 |
| hsa04722 | Neurotrophin signaling pathway | 0.50 | 0.73 | 4 |
| hsa04144 | Endocytosis |  | 1.00 | 3 |
| hsa04371 | Apelin signaling pathway | 1.00 | 1.00 | 3 |
| hsa04723 | Retrograde endocannabinoid signaling |  | 1.00 | 3 |
| hsa04727 | GABAergic synapse |  | 1.00 | 3 |
| hsa04014 | Ras signaling pathway | 0.83 | 0.86 | 3 |
| hsa04919 | Thyroid hormone signaling pathway | 0.75 | 0.86 | 3 |
| hsa04613 | Neutrophil extracellular trap formation | 0.44 | 0.75 | 3 |
| hsa04935 | Growth hormone synthesis, secretion and action |  | 0.75 | 3 |
| hsa04724 | Glutamatergic synapse |  | 1.00 | 2 |
| hsa04350 | TGF-beta signaling pathway | 0.88 | 0.80 | 2 |
| hsa04150 | mTOR signaling pathway |  | 0.80 | 2 |
| hsa04370 | VEGF signaling pathway | 0.75 | 0.80 | 2 |
| hsa04657 | IL-17 signaling pathway | 0.80 | 0.80 | 2 |
| hsa04916 | Melanogenesis |  | 0.80 | 2 |
| hsa04929 | GnRH secretion | 0.55 | 0.57 | 2 |
| hsa04080 | Neuroactive ligand-receptor interaction |  | 1.00 | 1 |
| hsa04668 | TNF signaling pathway | 1.00 | 1.00 | 1 |
| hsa04918 | Thyroid hormone synthesis |  | 1.00 | 1 |

For each neighborhood, one through five, surrounding angiotensin/renin (yellow, RAS) nodes, pathways through edges were examined for dopamine/dopaminergic (magenta, DA) node proximity. If the path contained only DA associated (cyan, DA_assoc_) and/or RAS associated (gray, RAS_assoc_) nodes, the yellow node was marked with a blue cross (true positive, TP) for ‘in’ or a cyan cross (false negative, FN) for ‘out’ of the neighborhood being examined. If the contiguous path contained nodes other than RAS_assoc_ or DA_assoc_ (pale green), the yellow node was marked with a black cross (false positive, FP) for ‘in’ or a gray cross (true negative, TN) for ‘out’ of the neighborhood being examined (Figure 3).

**Figure 3.**
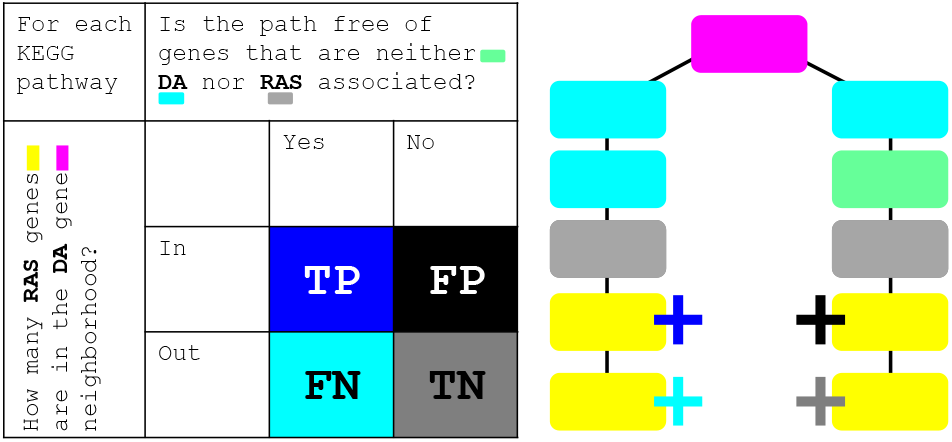
Contingency test and fourth neighborhood example.

All calculations were performed in Excel (Microsoft Corp., WA), ImageJ (Rasband, 1997-2018) was used for tabulation of supplemental gif images and Cytoscape (Shannon et al., 2003) was used for plotting the network in figure 4.

**Figure 4.**
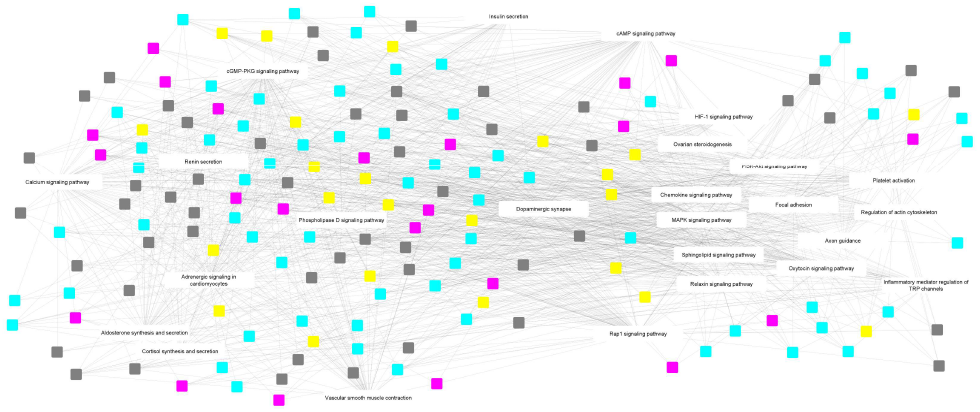
Naive unstructured network of 25 (F-AUC)_0._85-5 pathways and 160 four KP proteins [RAS|RAS_assoc_|DA|DA_assoc_].

## RESULTS

The 3309 proteins from the 20 Swiss-Prot keyword searches are in Supplementary Table 1. For each entry, membership in a keyword search is tabulated as a ‘1’. This data was used to calculate the binary count clustering for Swiss-Prot keyword searches and visualize the membership maps in Figure 2.

The contingency testing shown in the 51 supplemental gif images as referenced in Table 1 [1^st^, 2^nd^, 3^rd^, 4^th^, 5^th^ neighborhoods, unmasked KEGG output] is a graphical representation of the exact F-AUC ROC analysis procedures.

The 51 KEGG pathways in Table 1 are color coded for all F-AUC-3 [specifically (F-AUC)_0.7_-3, maximum AUC or F-measure at least 0.7 and 3 TP nodes (Westwood et al., 2022)] blue for F-AUC of 1, orange for F-AUC at least 0.85 and gray for F-AUC at least 0.7. Of the 25 (F-AUC)_0.85_-5 pathways (bold), there were 160 proteins in at least 4 KEGG pathways (KP). There were 23 RAS (7.7±0.72 KP, mean+sem), 51 RAS_assoc_ (7.5±0.59 KP), 22 DA (7.6±0.98 KP), 64 DA_assoc_ (7.3±0.44 KP) proteins (Chi-square, p=0.44, ns; Figure 4).

## DISCUSSION

This framework ties the RAS and DA systems using targeted Swiss-Prot curated database search output membership in KEGG networks by implementing CPDB, resulting in an enriched dataset with a balanced representation from RAS, RAS_assoc_, DA and DA_assoc_ proteins for further analysis. The utility if this method is in the simplicity of the contingency testing on display in the supplemental gif images. The ontological microenvironment defined by this framework array is broadly applicable to describing biological systems. This Swiss-Prot/KEGG/CPDB conduit can provide foundational information in the investigation of novel networks.

## ACKNOWLEDGMENT

Supported by WFSM CVSC Pilot Award, CTSI NCATS UL1TR001420 and Odell Farley Foundation.

## ABBREVIATIONS

DCS: discrete correlate summation
KEGG: Kyoto Encyclopedia of Genes and Genomes
CPDB: ConsensusPathDB-human
RL-OriMaTArA: original matrix total array average was calculated right to left
AUC: Area under the curve
ROC: receiver operating characteristic
RAS: angiotensin/renin
DA: dopamine/dopaminergic
DA_assoc_: DA associated
RAS_assoc_: RAS associated
TP: true positive
FN: false negative
FP: false positive
TN: true negative

## REFERENCES

Kanehisa M, Goto S. KEGG: kyoto encyclopedia of genes and genomes. Nucleic Acids Res. 2000 Jan 1;28(1):27–30. doi: 10.1093/nar/28.1.27. PMID: 10592173; PMCID: PMC102409.

Kamburov A, Pentchev K, Galicka H, Wierling C, Lehrach H, Herwig R. ConsensusPathDB: toward a more complete picture of cell biology. Nucleic Acids Res. 2011 Jan;39(Database issue):D712–7. doi: 10.1093/nar/gkq1156. Epub 2010 Nov 11. PMID: 21071422; PMCID: PMC3013724.

The UniProt Consortium, UniProt: the universal protein knowledgebase in 2021, Nucleic Acids Research, Volume 49, Issue D1, 8 January 2021, Pages D480–D489, 10.1093/nar/gkaa1100

Bronson S, Westwood B, Cook K, Emenaker N, Chappell M, Roberts D, Soto-Pantoja D. Discrete Correlation Summation Clustering Reveals Differential Regulation of Liver Metabolism by Thrombospondin-1 in Low-Fat and High-Fat Diet-Fed Mice. Metabolites 2022, 12, 1036. 10.3390/metabo12111036

Fawcett T. An introduction to ROC analysis. Pattern Recogn Lett. 2006;27(8):861–74.

Rasband, W.S., ImageJ, U. S. National Institutes of Health, Bethesda, Maryland, USA, https://imagej.net/ij/, 1997-2018.

Shannon P, Markiel A, Ozier O, Baliga NS, Wang JT, Ramage D, Amin N, Schwikowski B, Ideker T. Cytoscape: a software environment for integrated models of biomolecular interaction networks. Genome Research 2003 Nov; 13(11):2498–504

Westwood B, Patil P, Tallant A, Gallagher P. Microbiome Metabolic Network Expansion Following Muscadine Grape Extract Intervention of Hypertensive Rats SB3C2022–266, 2022. https://archive.sb3c.org/sb3c2022/

