## Supplementary figures and images for "Discrete Correlate Summation Clustering of a Dopamine-Angiotensin Network"

### 4010

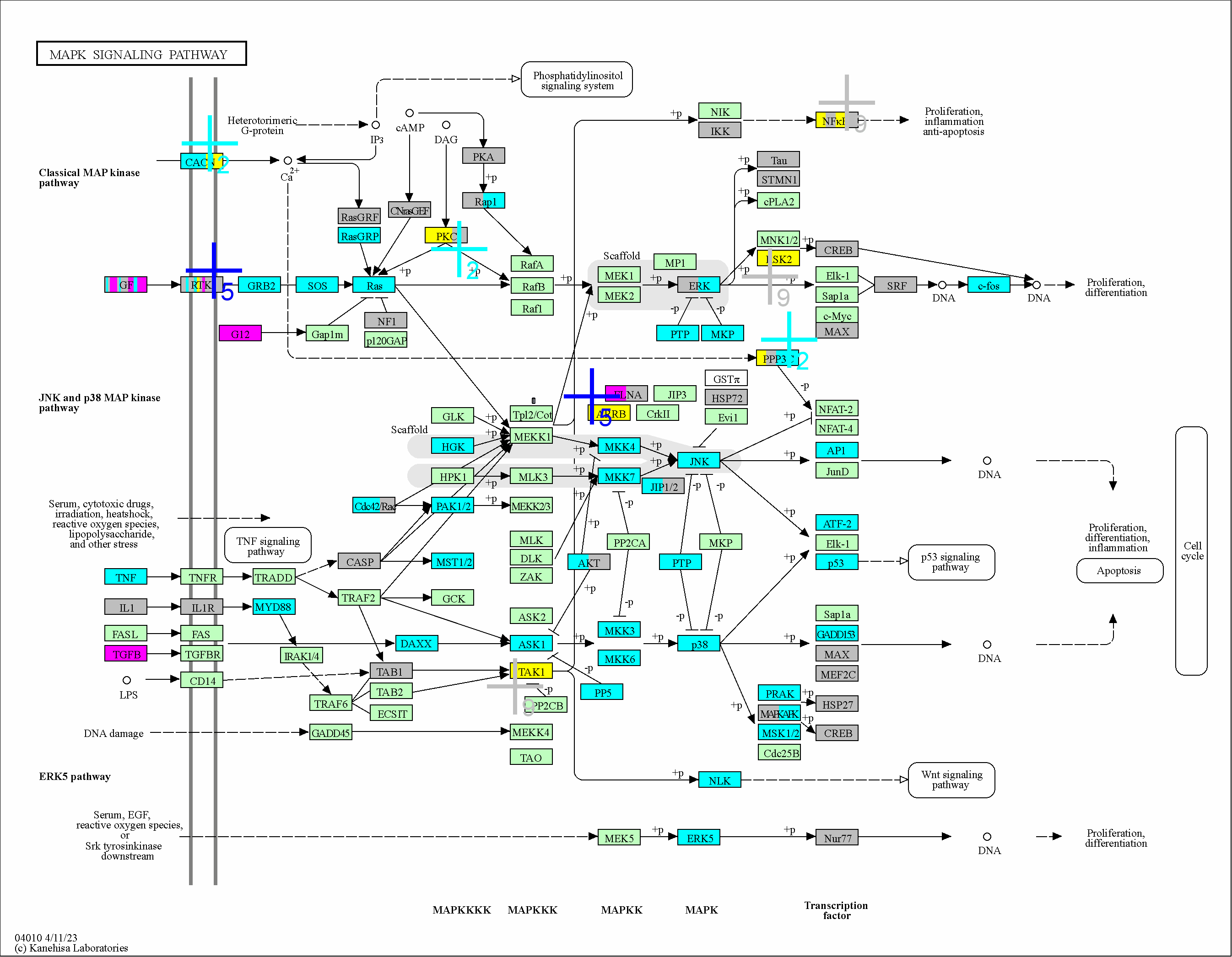

### 4014

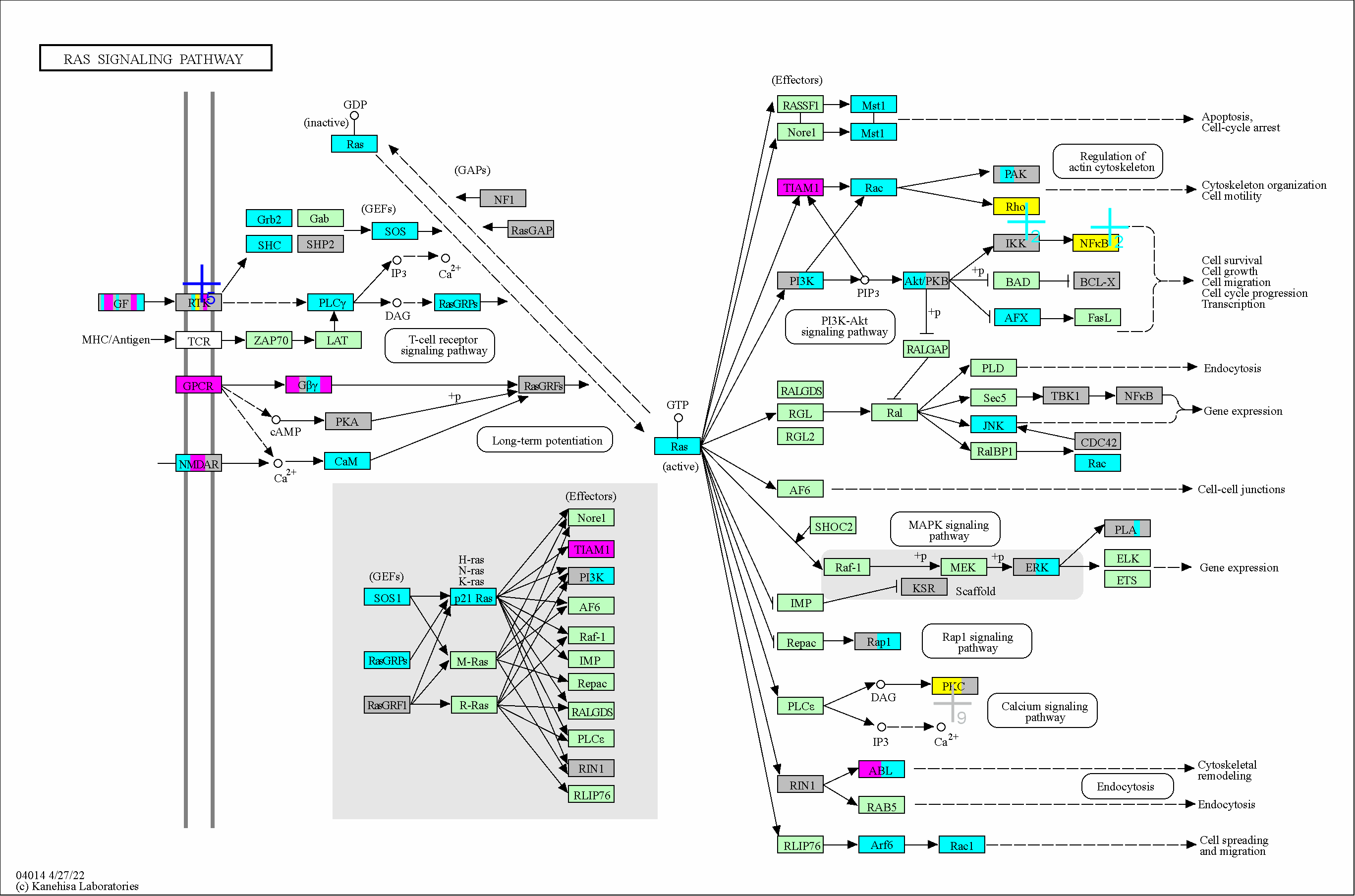

### 4015

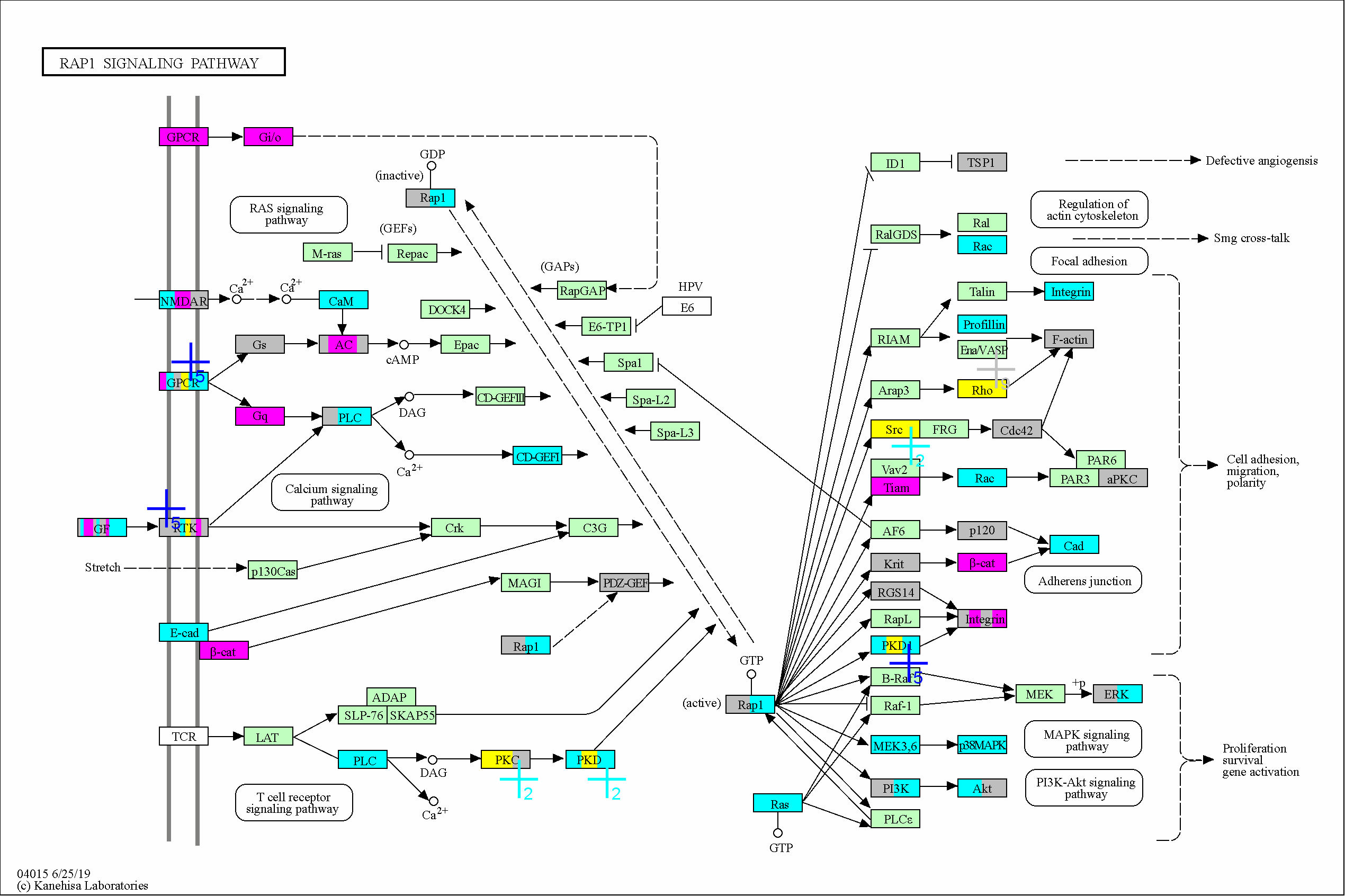

### 4020

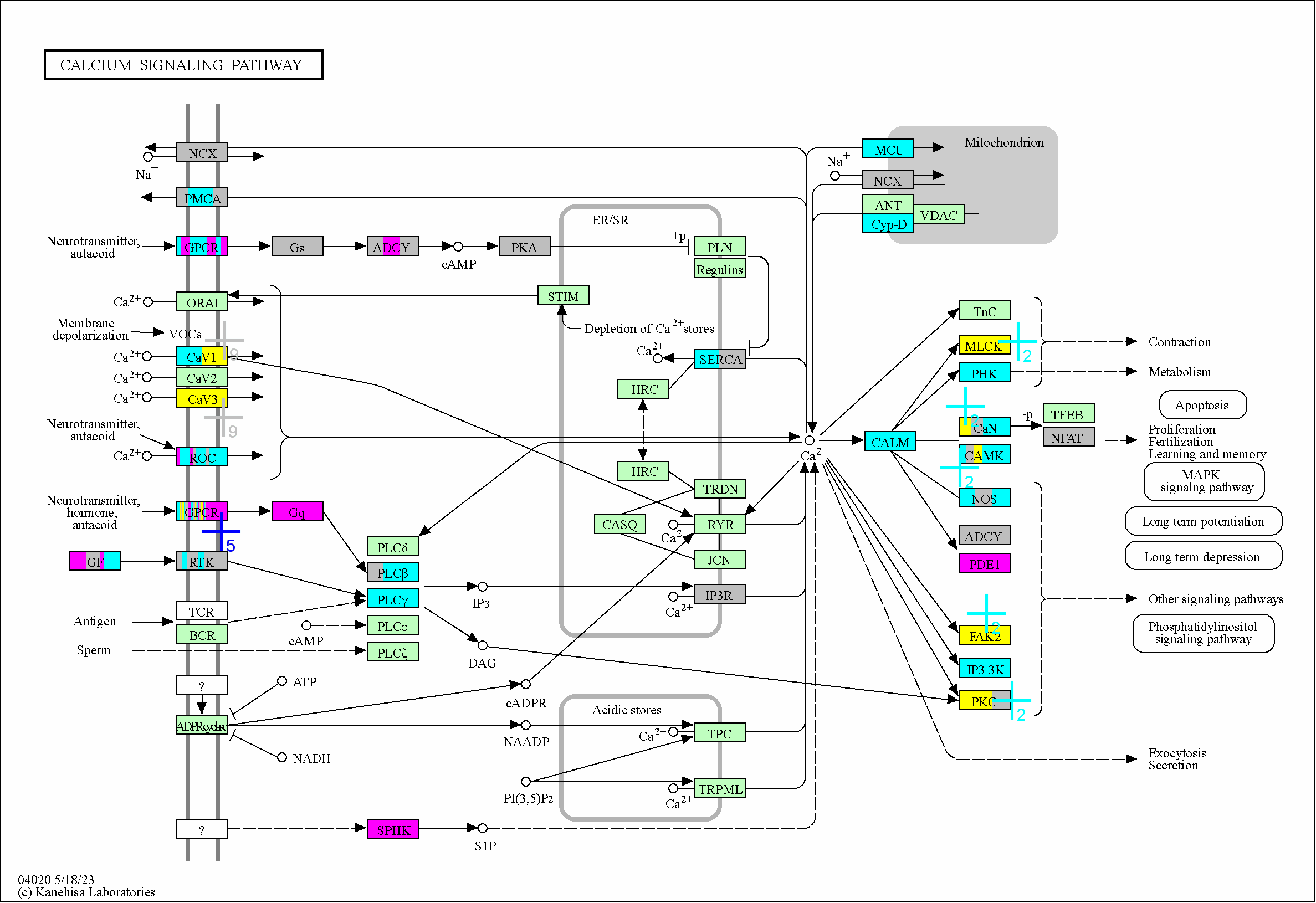

### 4022

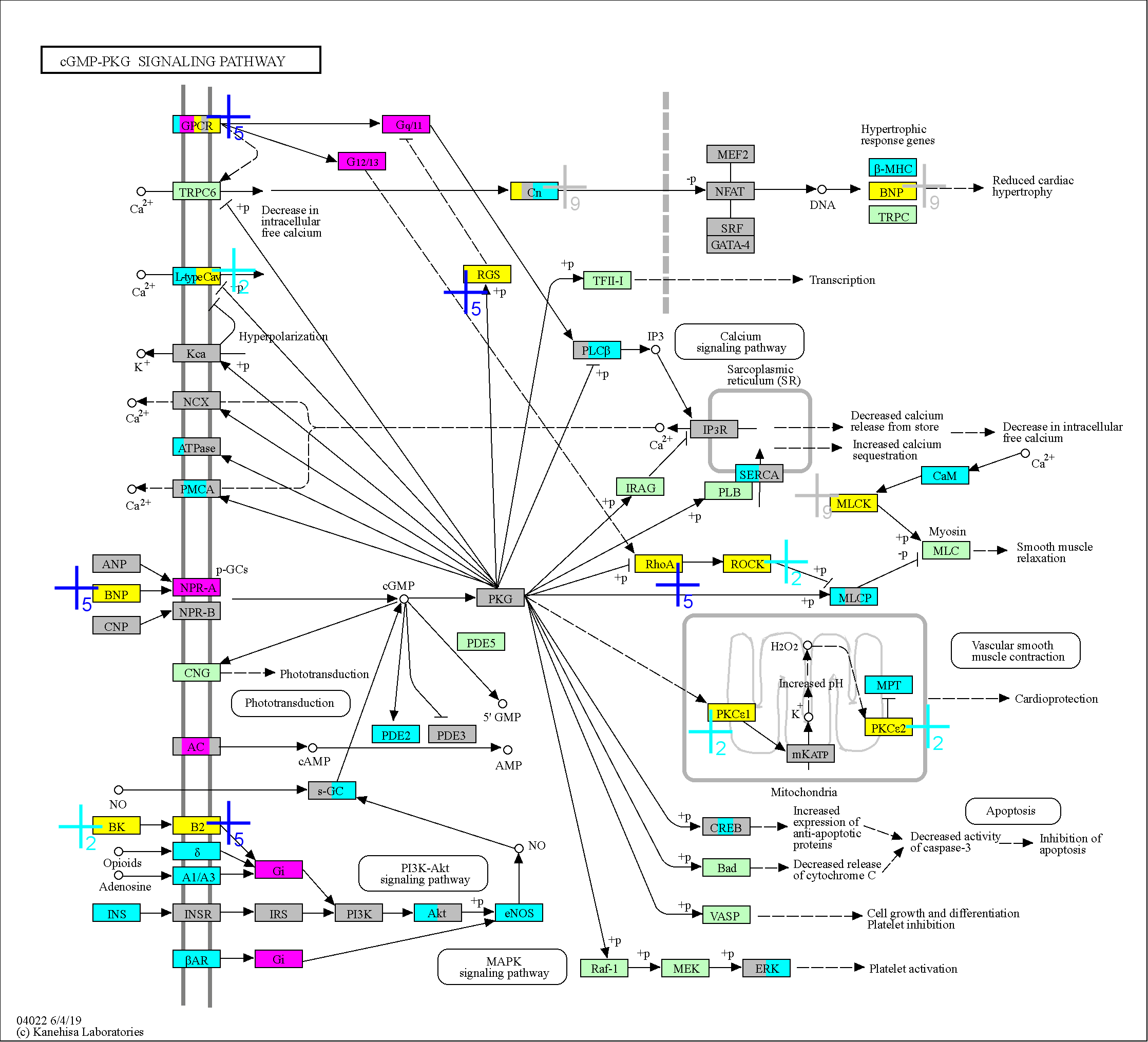

### 4024

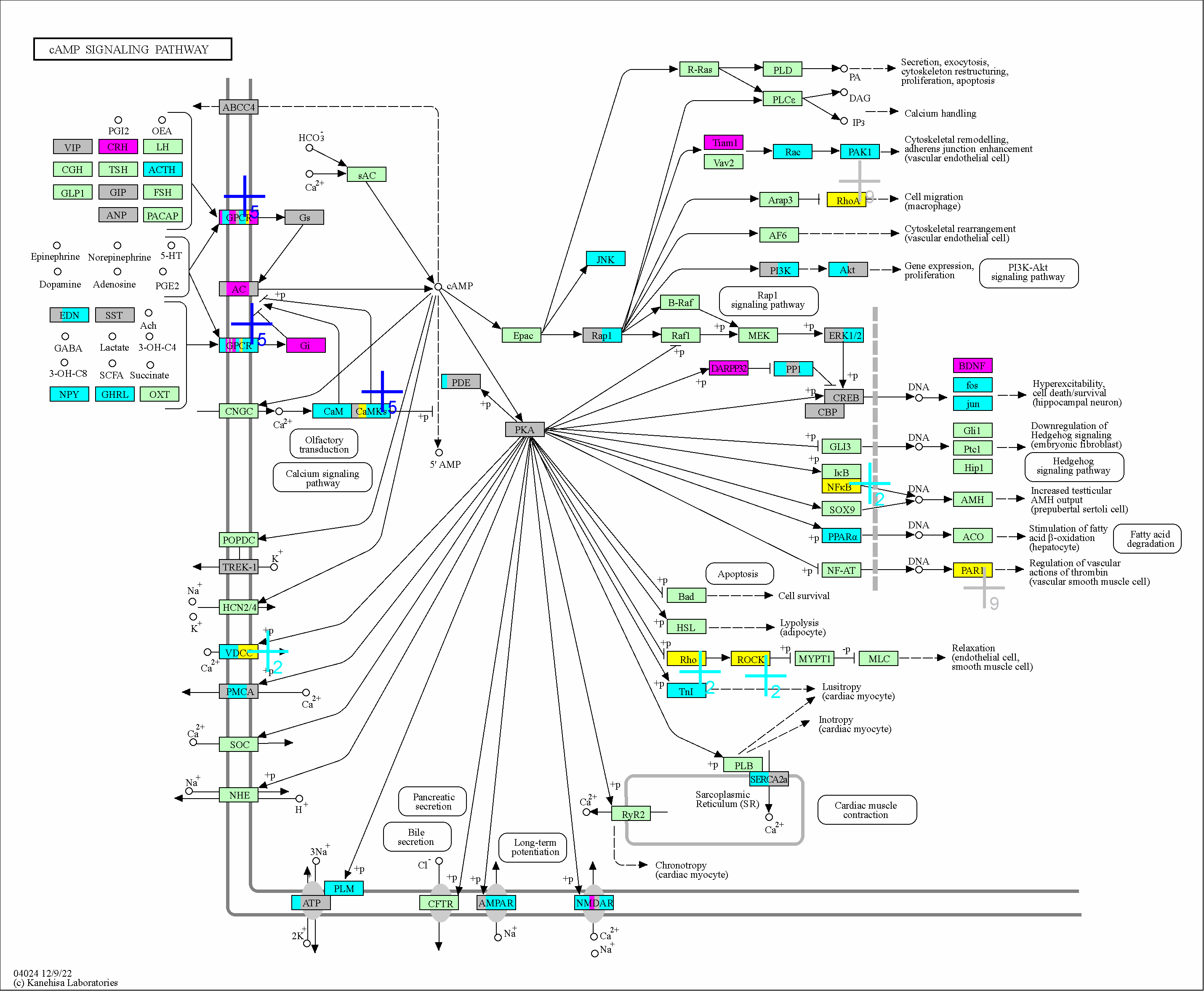

### 4062

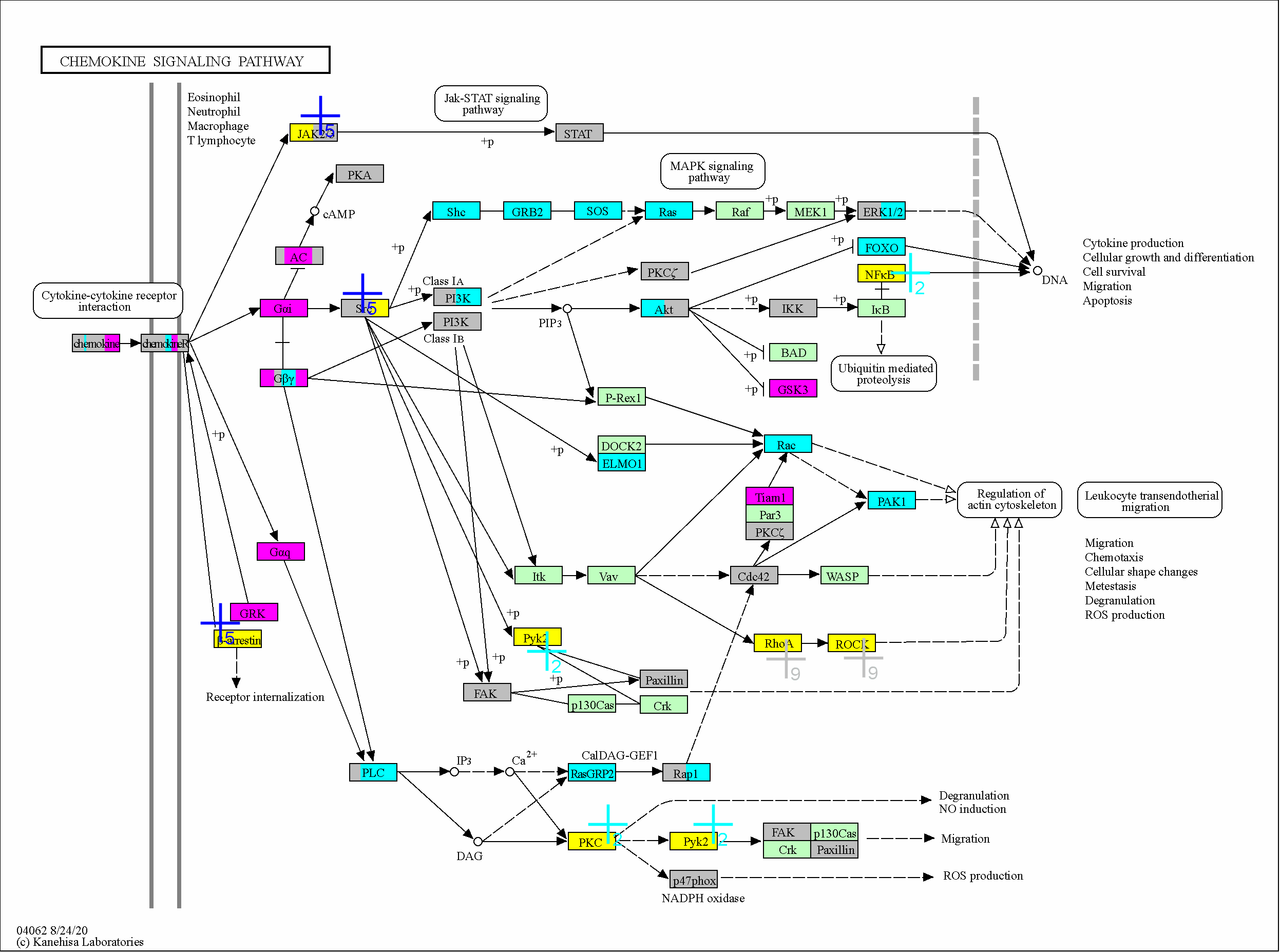

### 4066

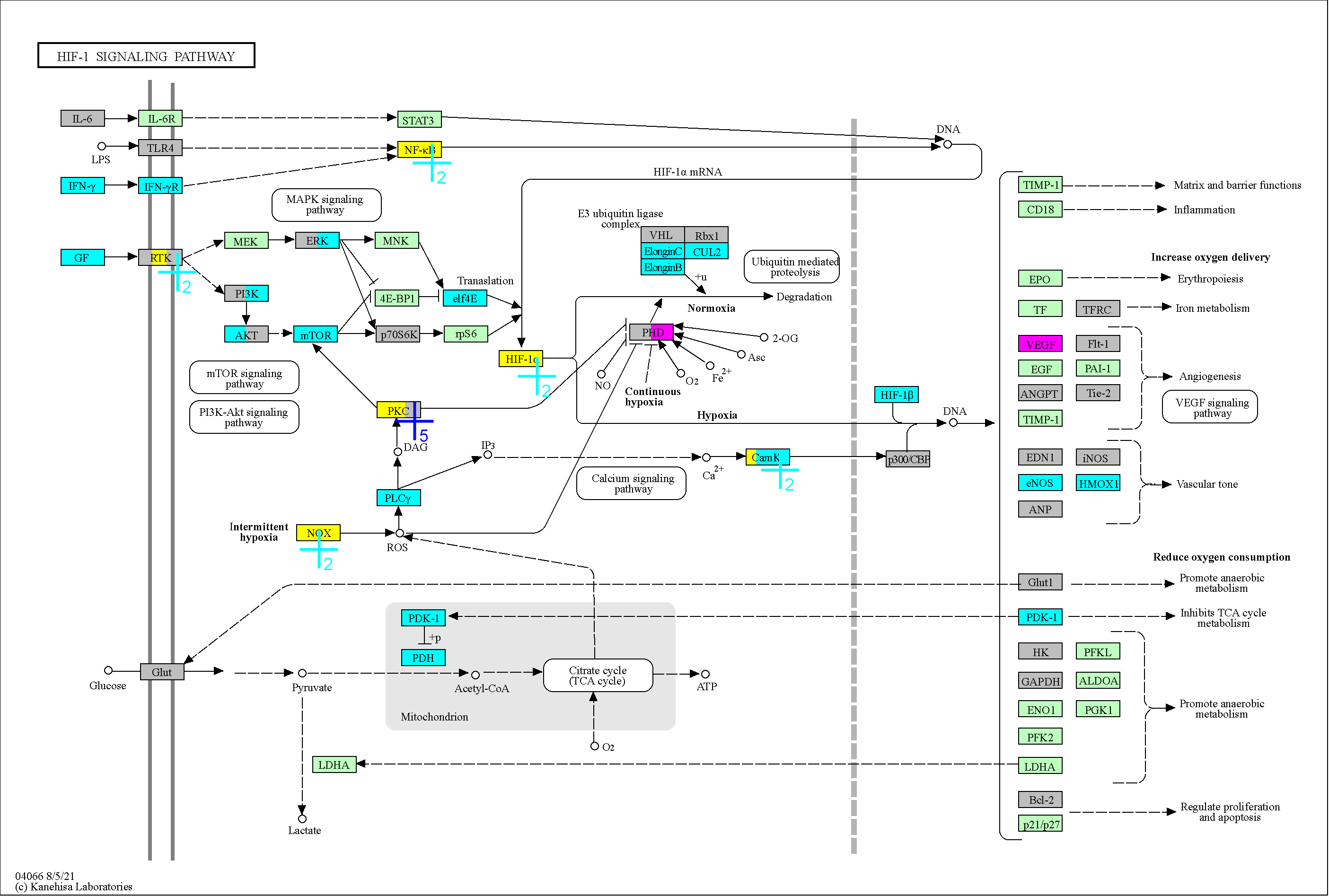

### 4071

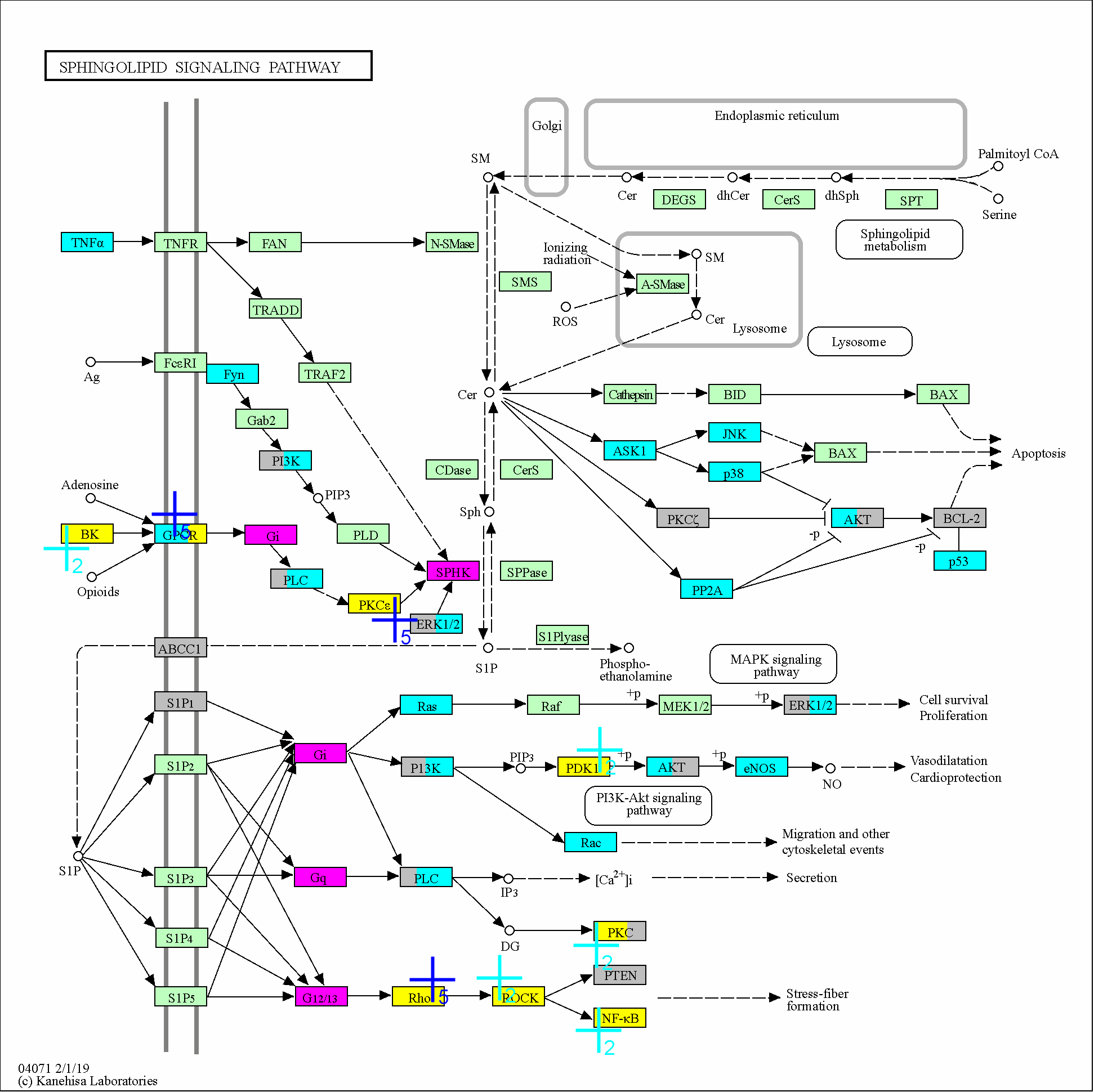

### 4072

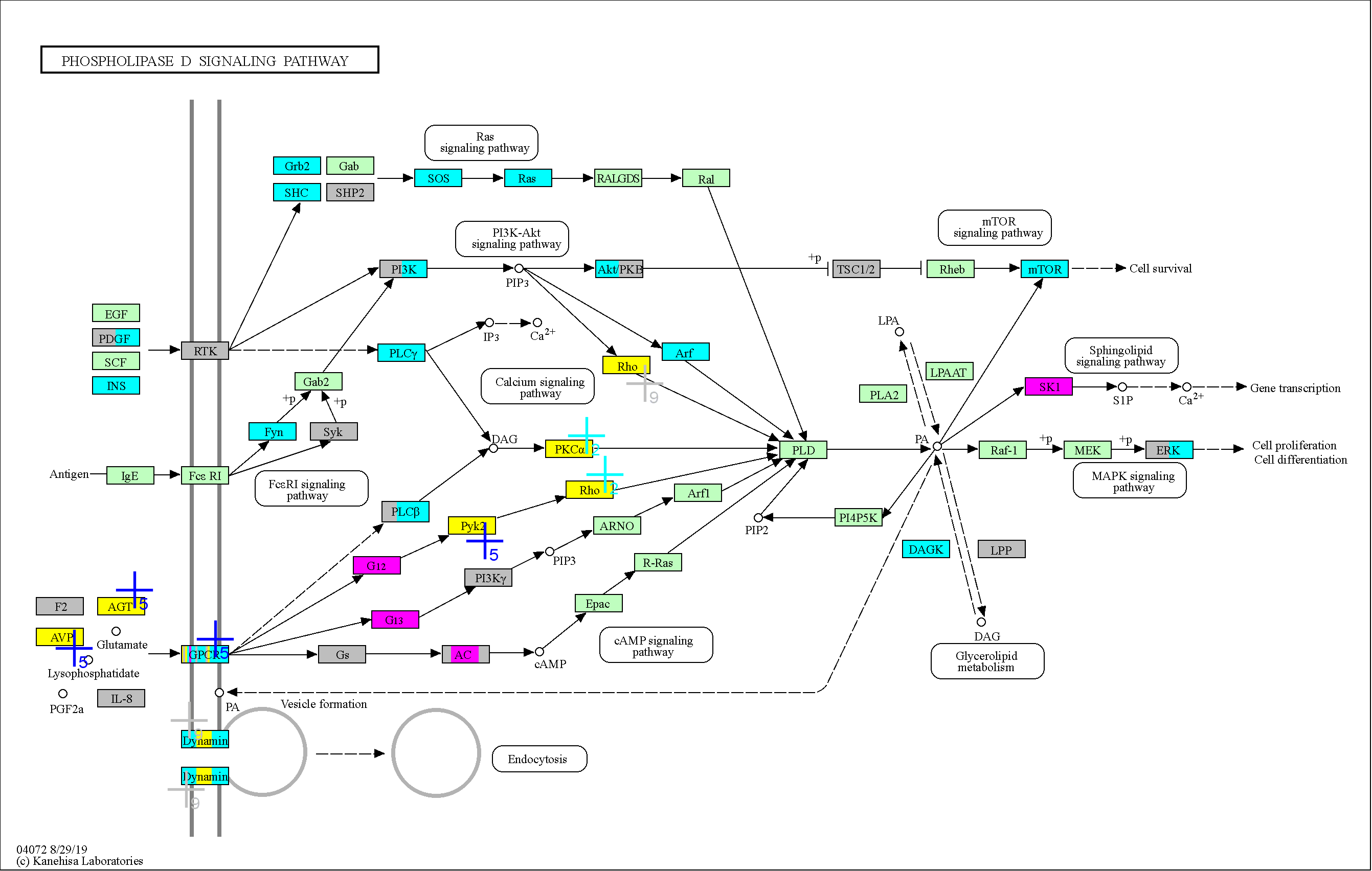

### 4080

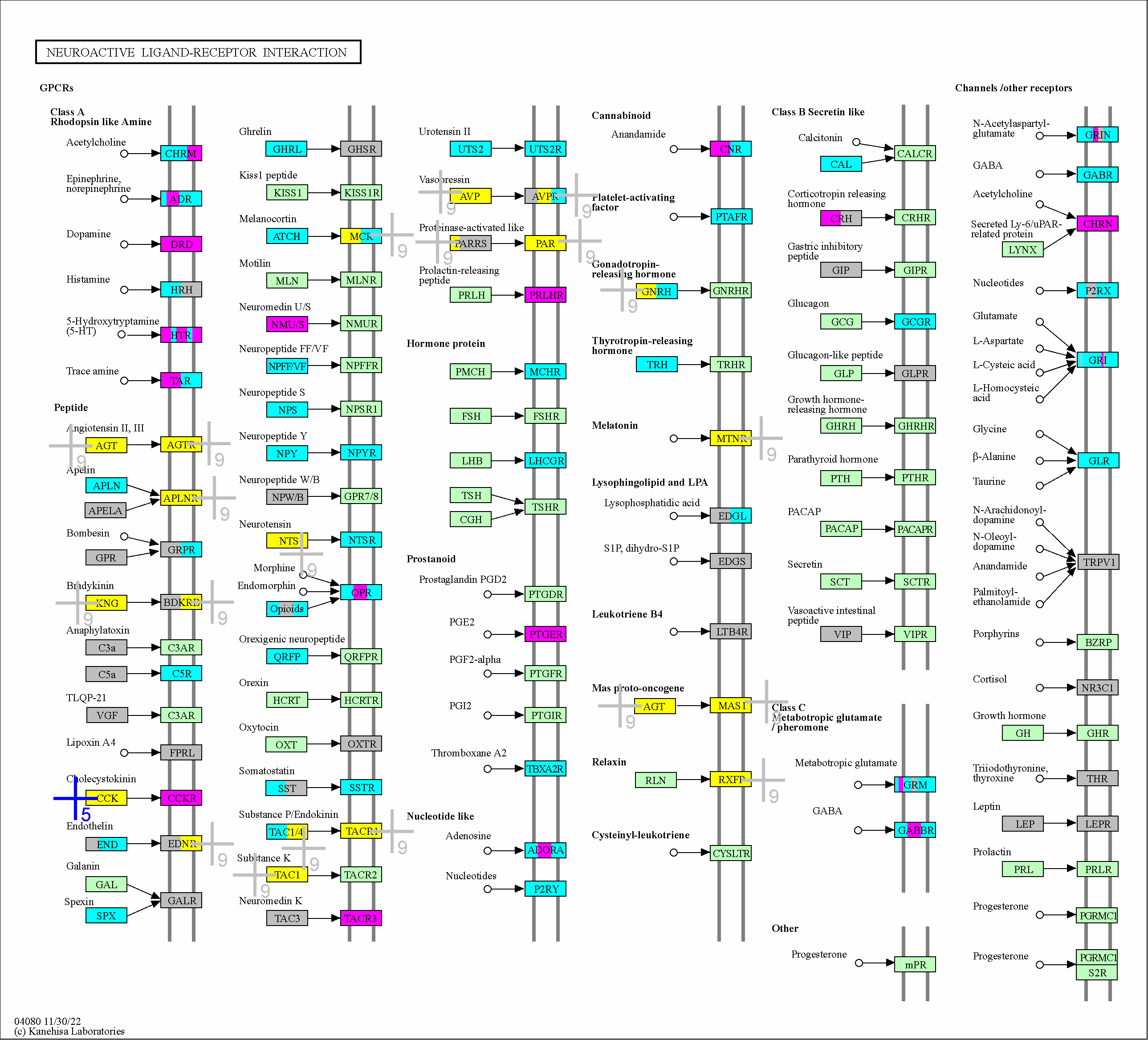

### 4144

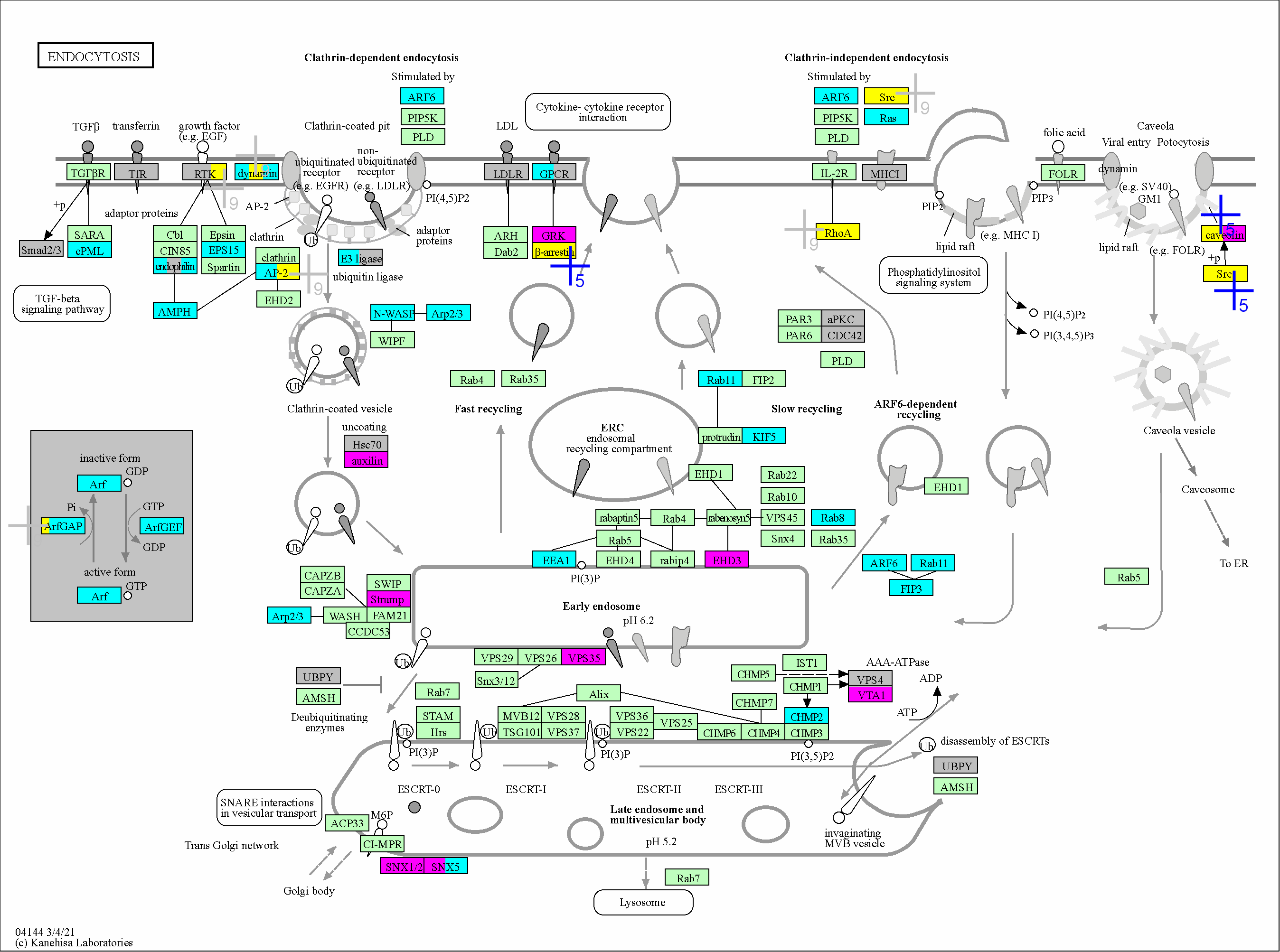

### 4150

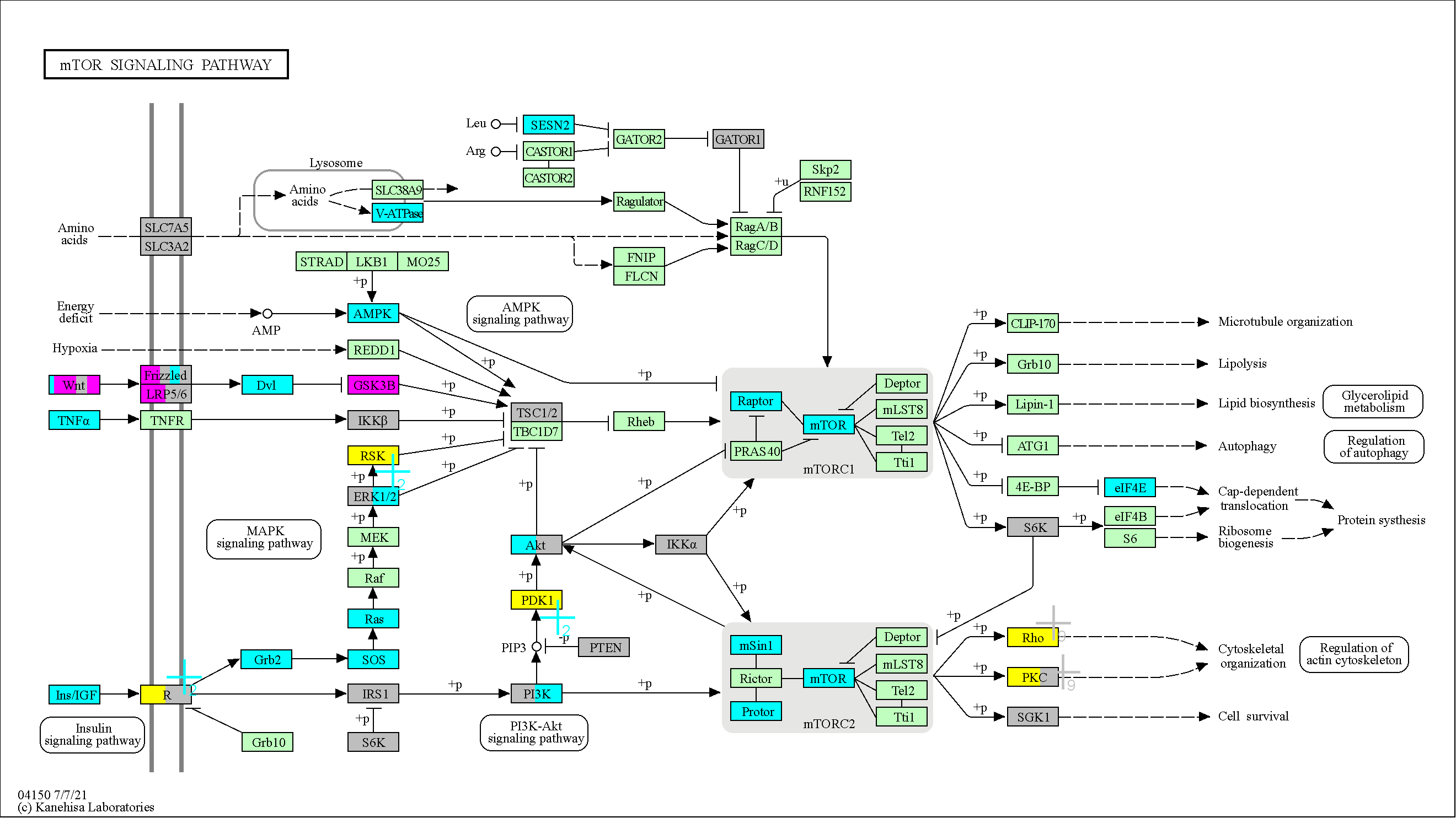

### 4151

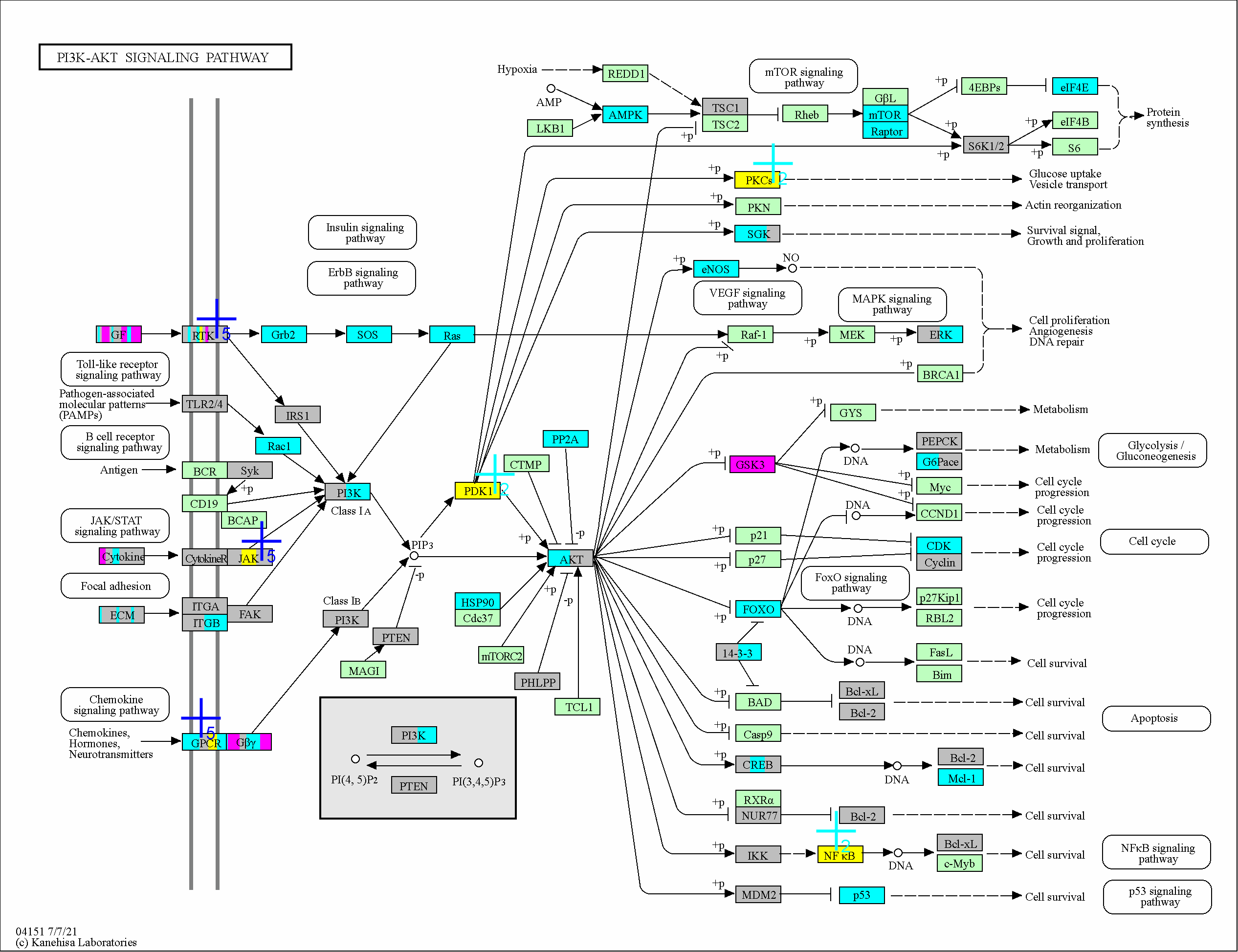

### 4261

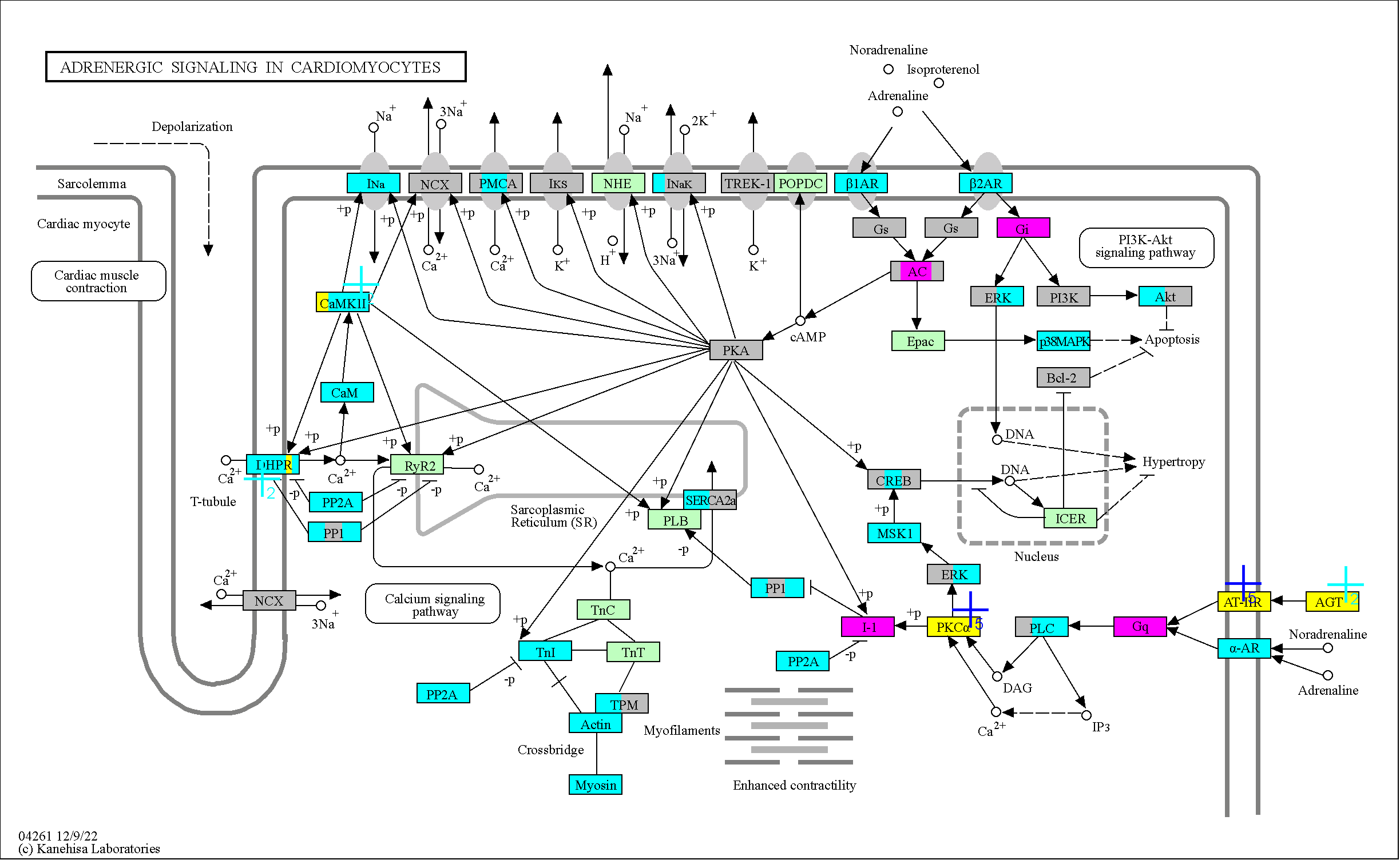

### 4270

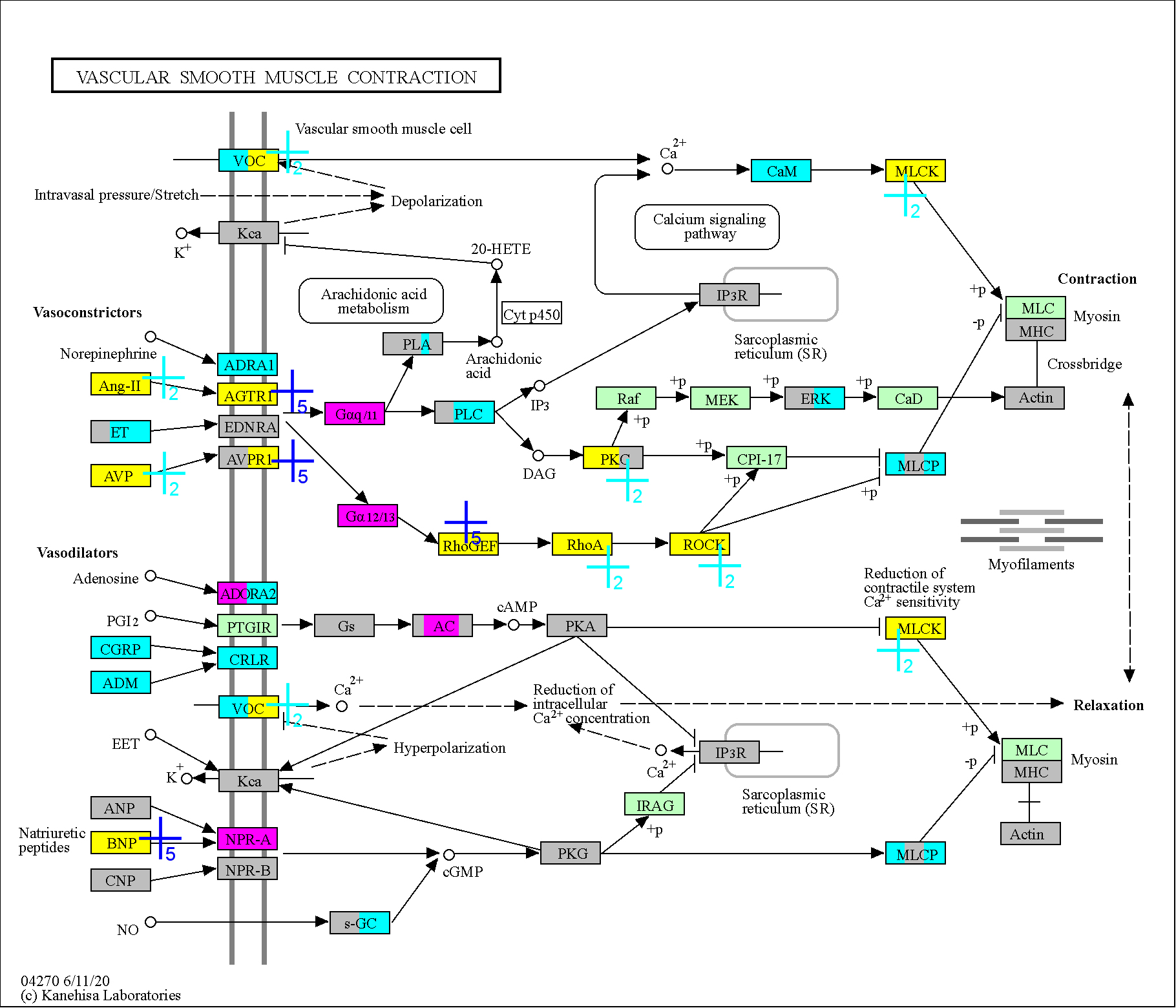

### 4310

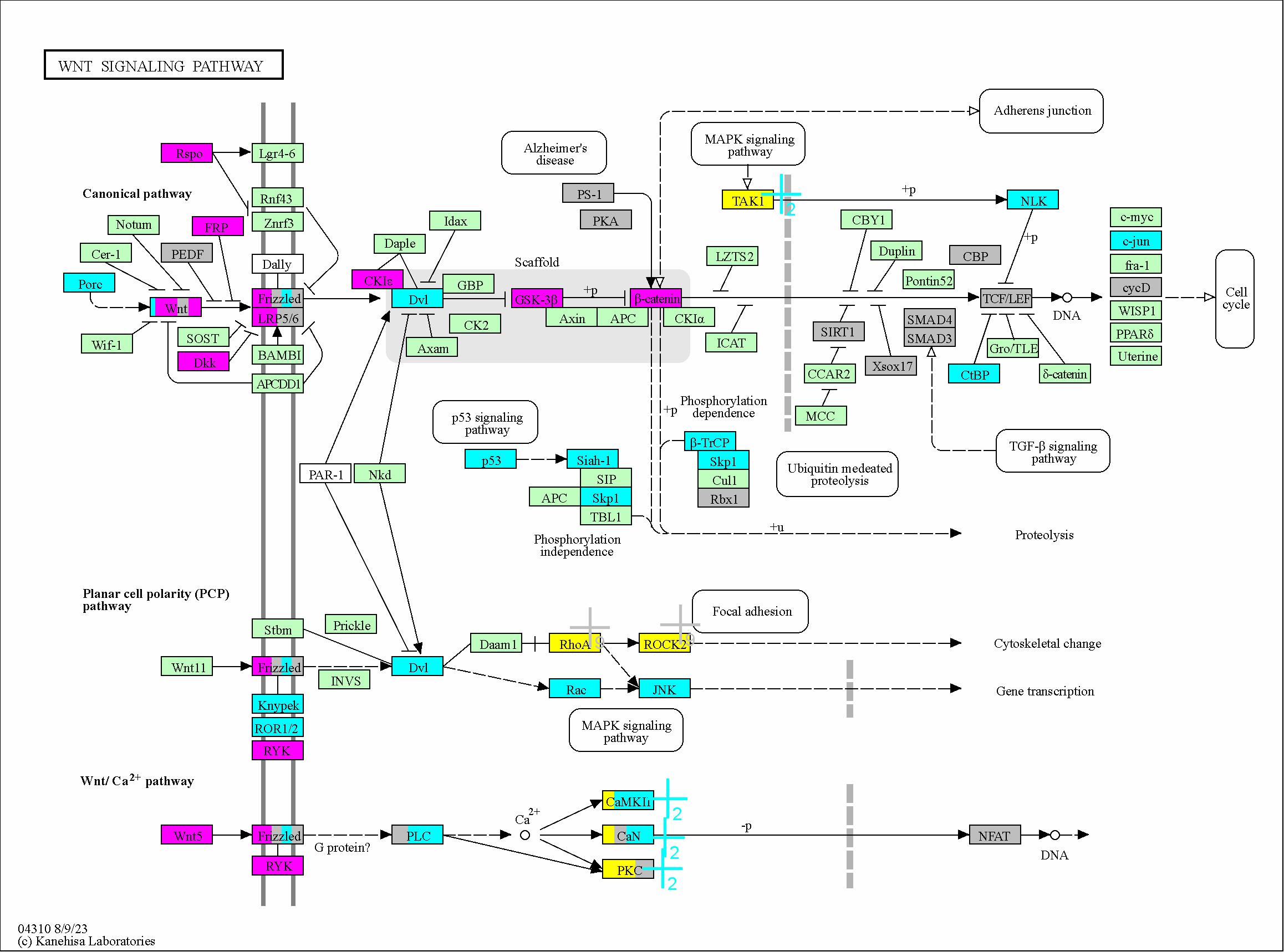

### 4350

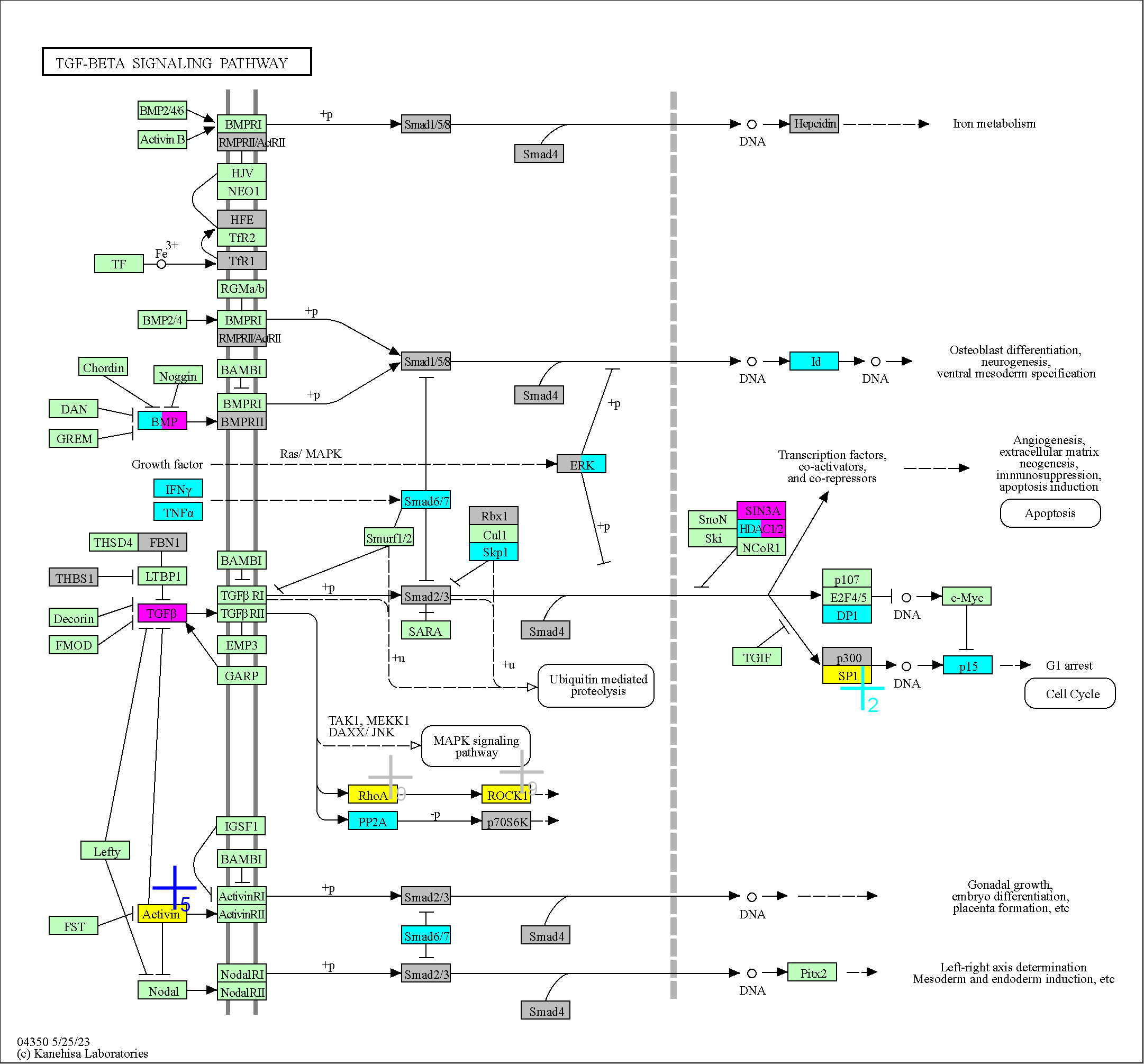

### 4360

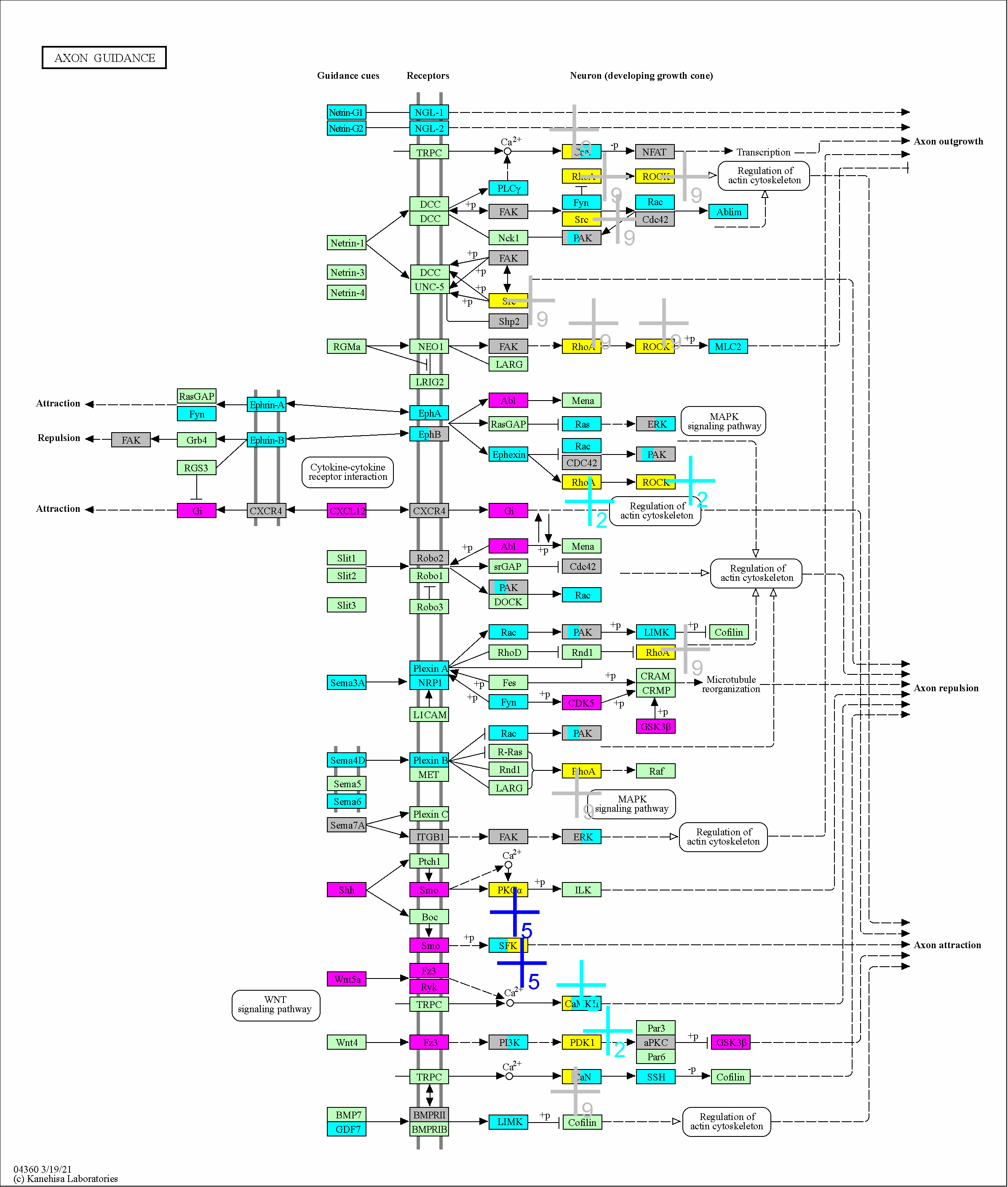

### 4370

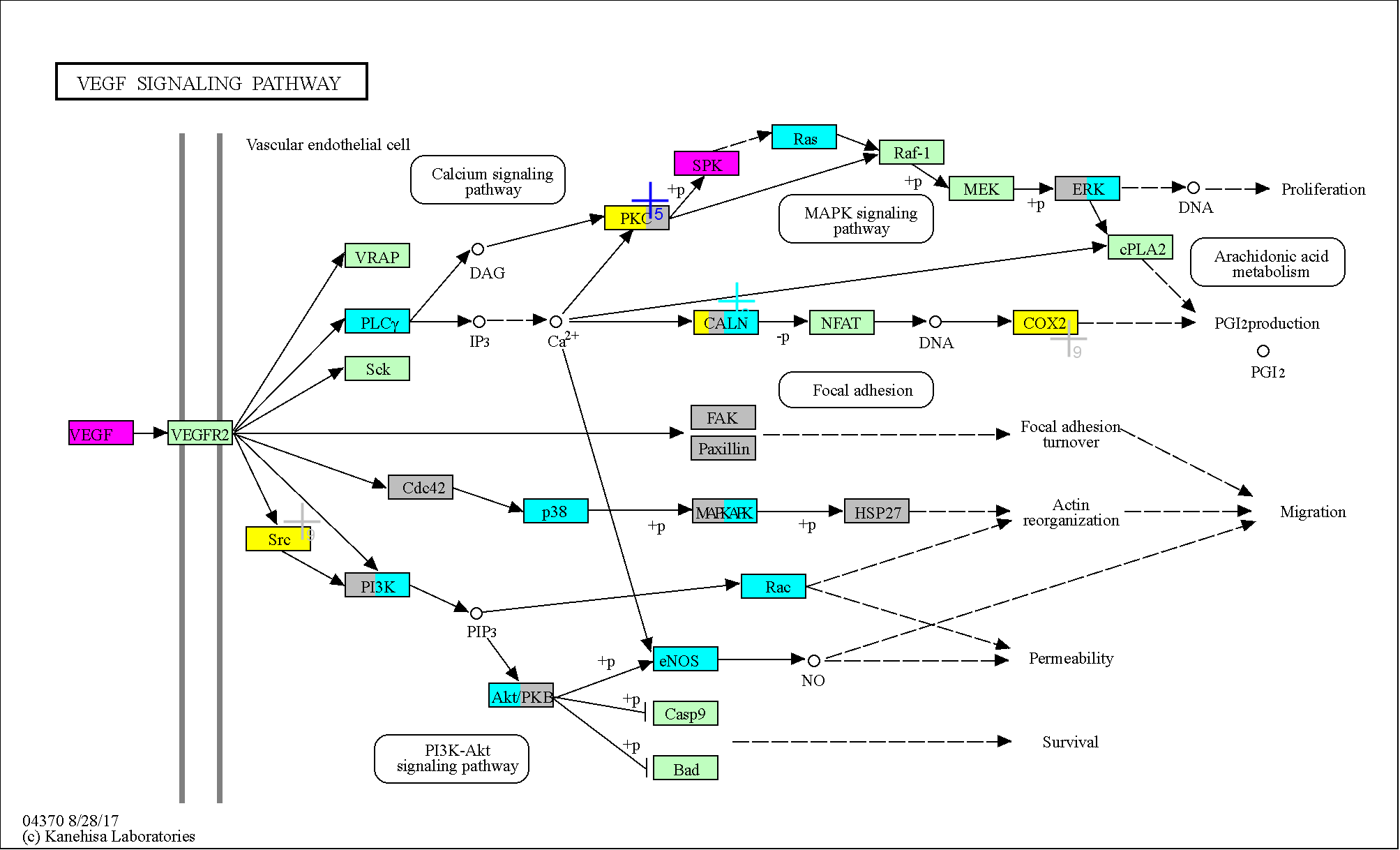

### 4371

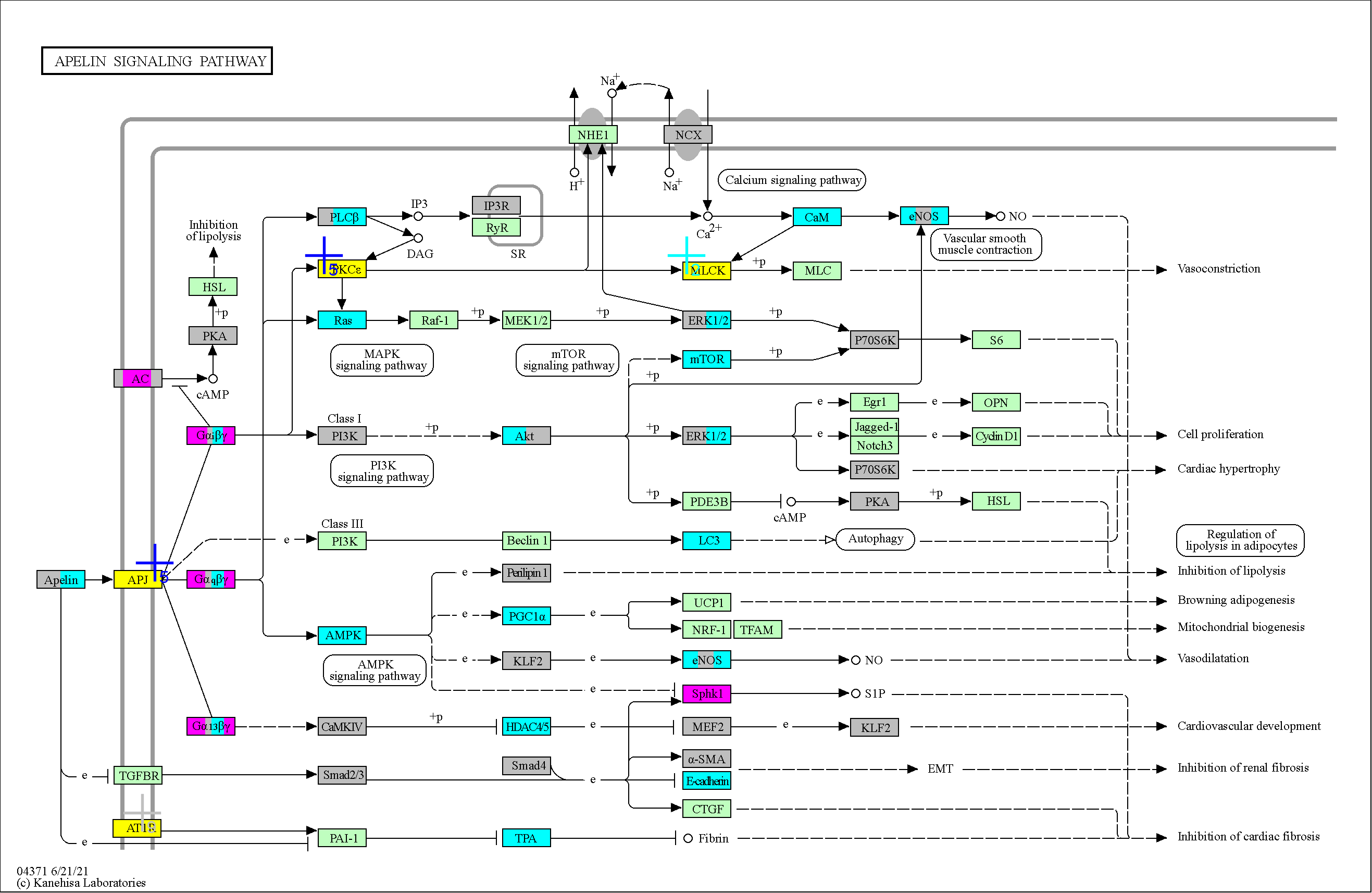

### 4510

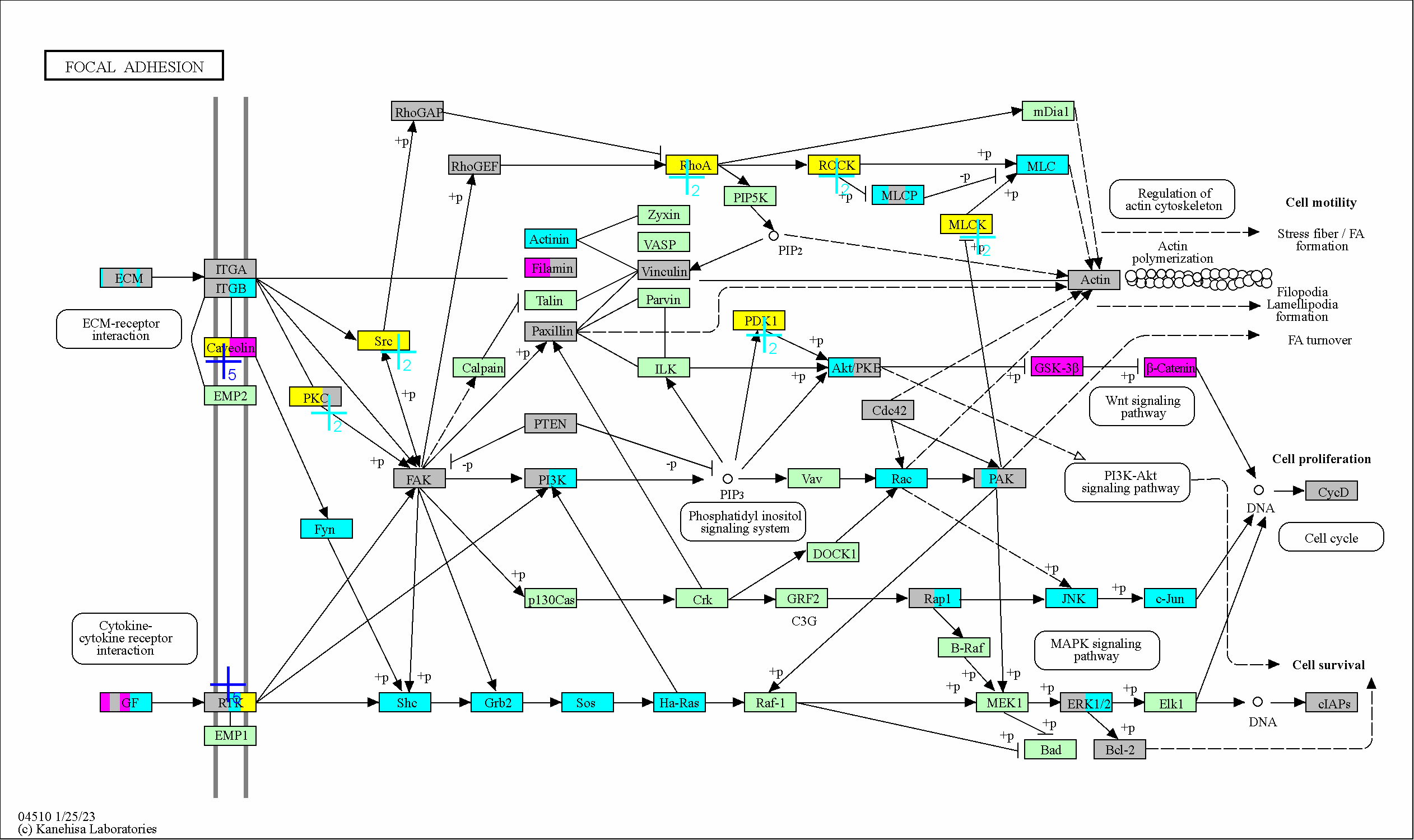

### 4611

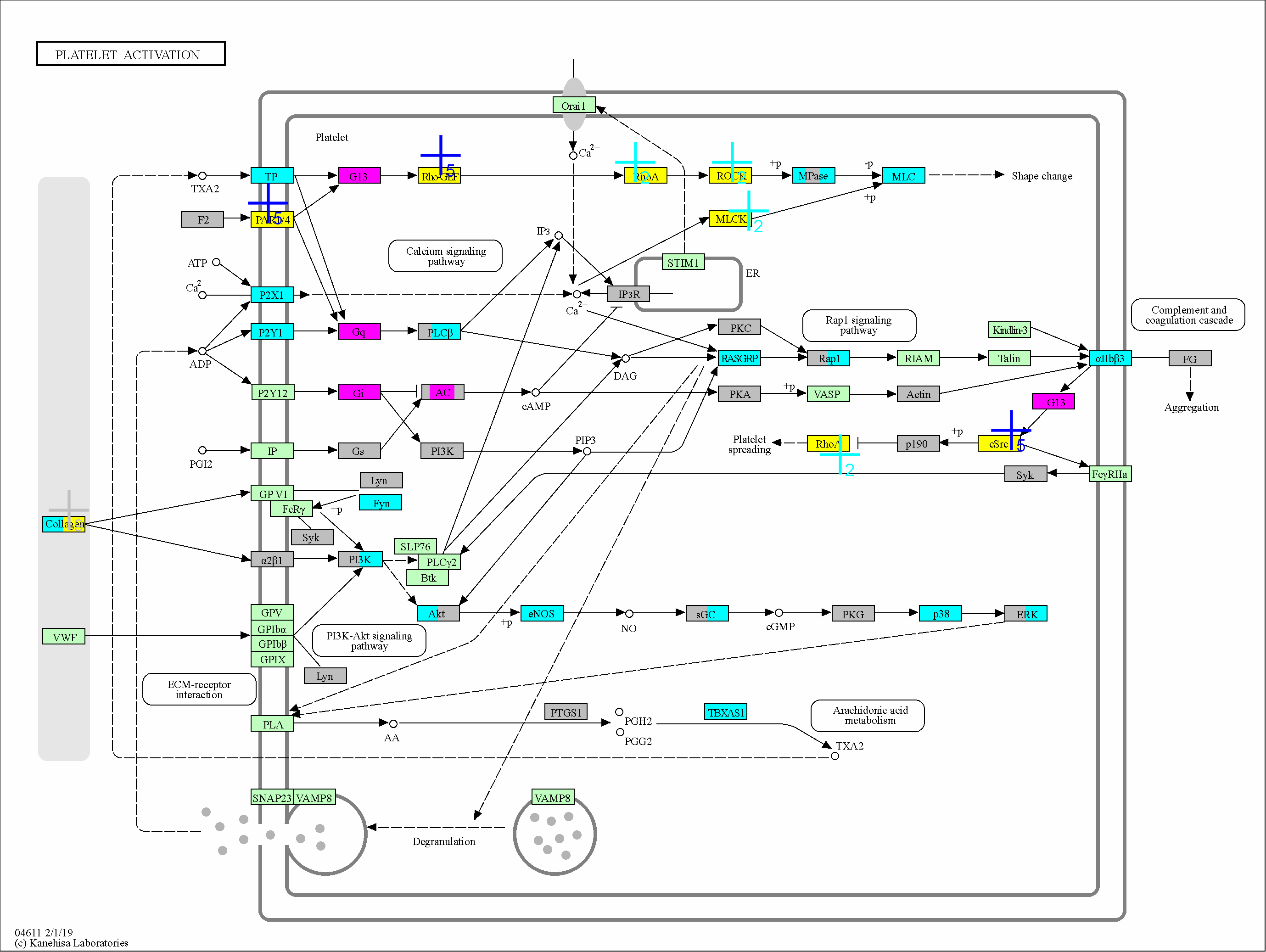

### 4613

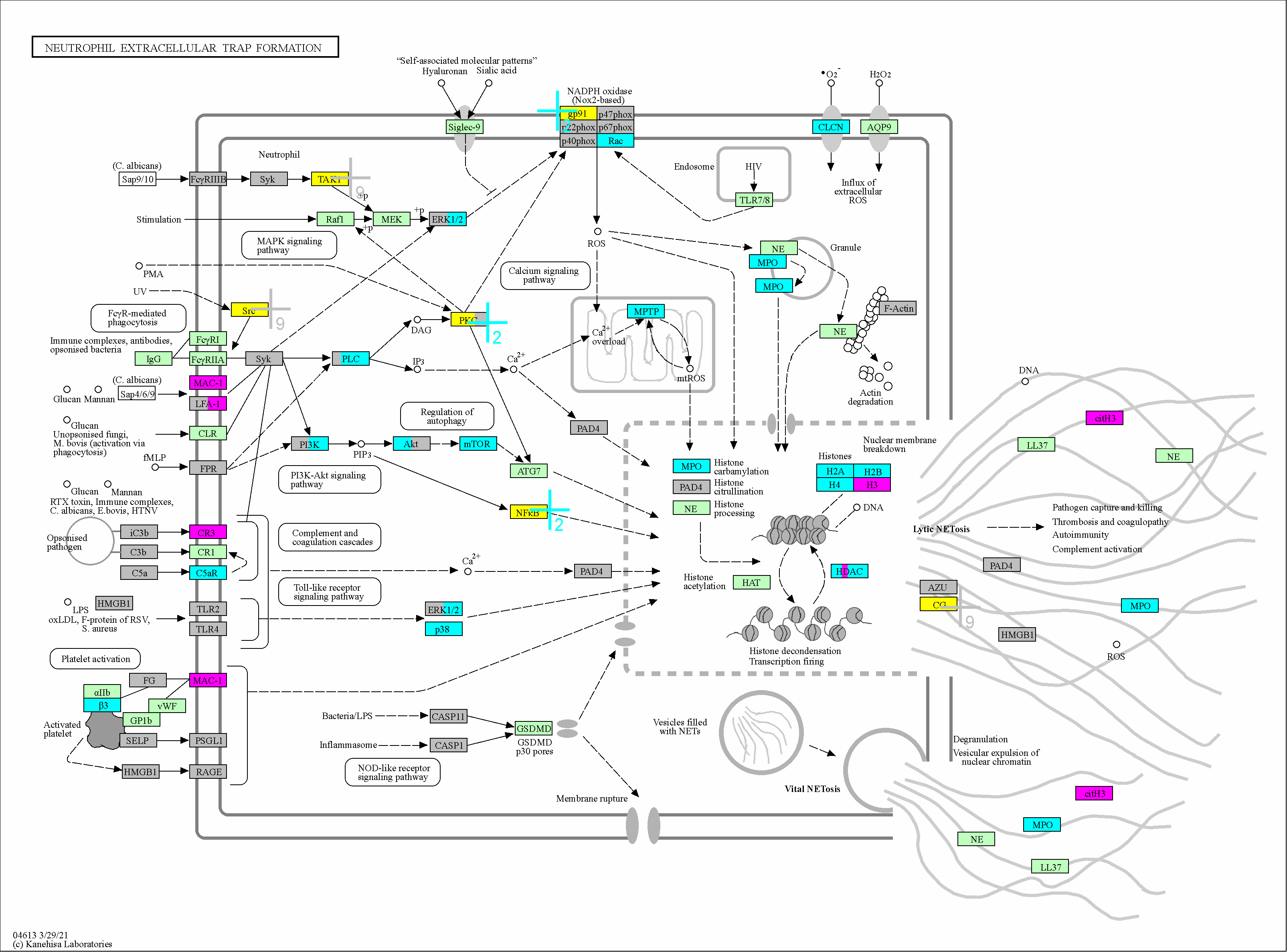

### 4657

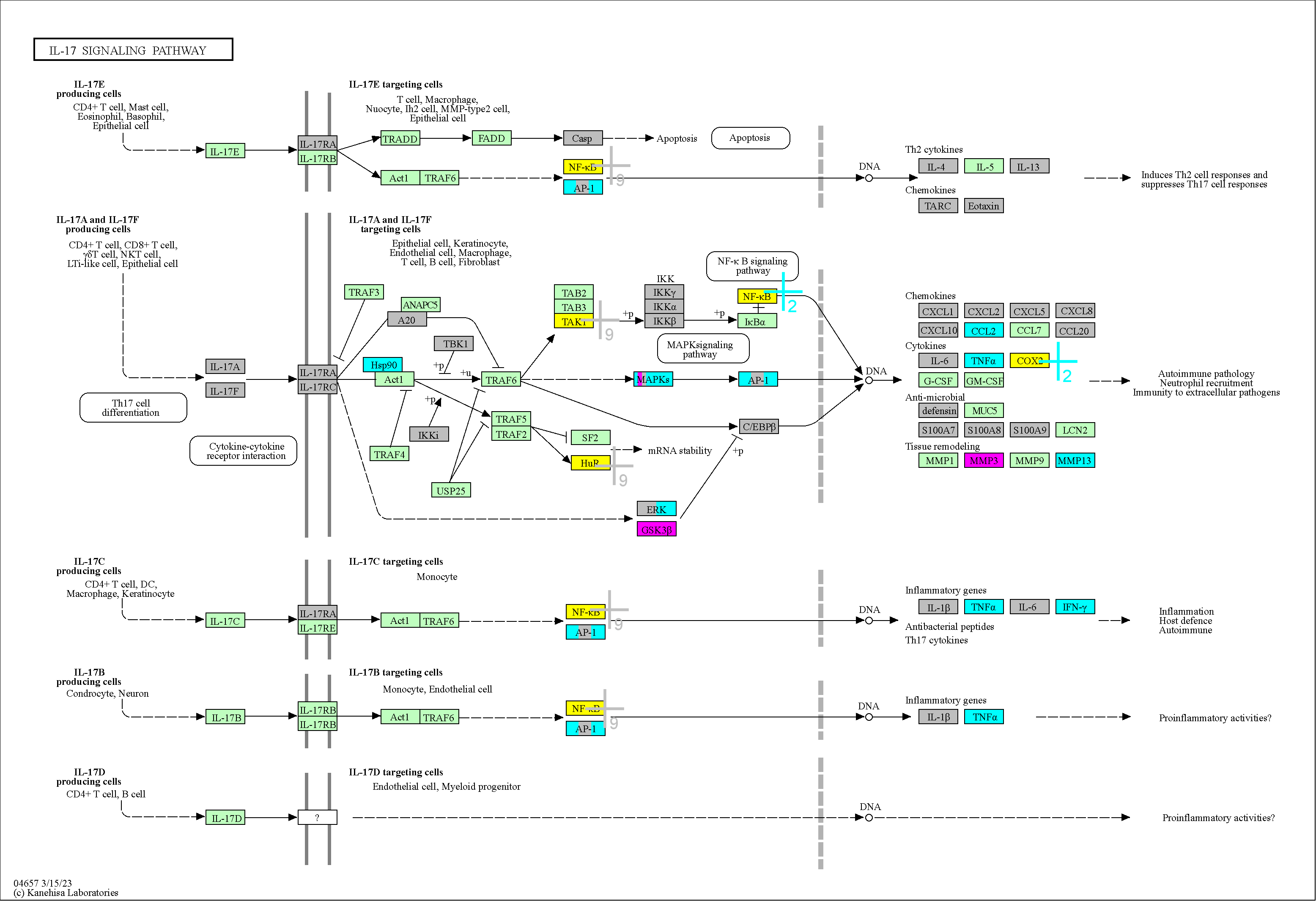

### 4668

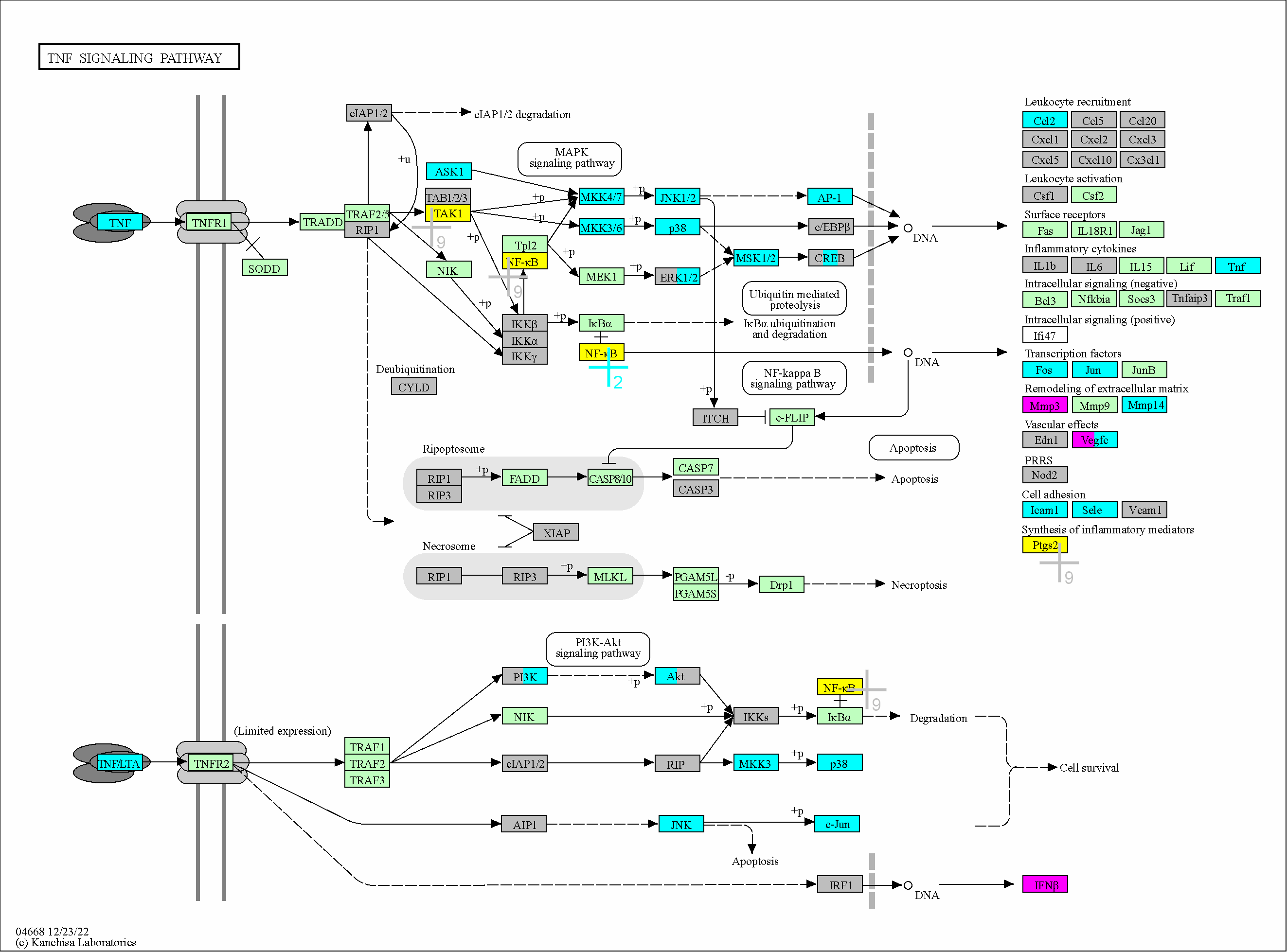

### 4670

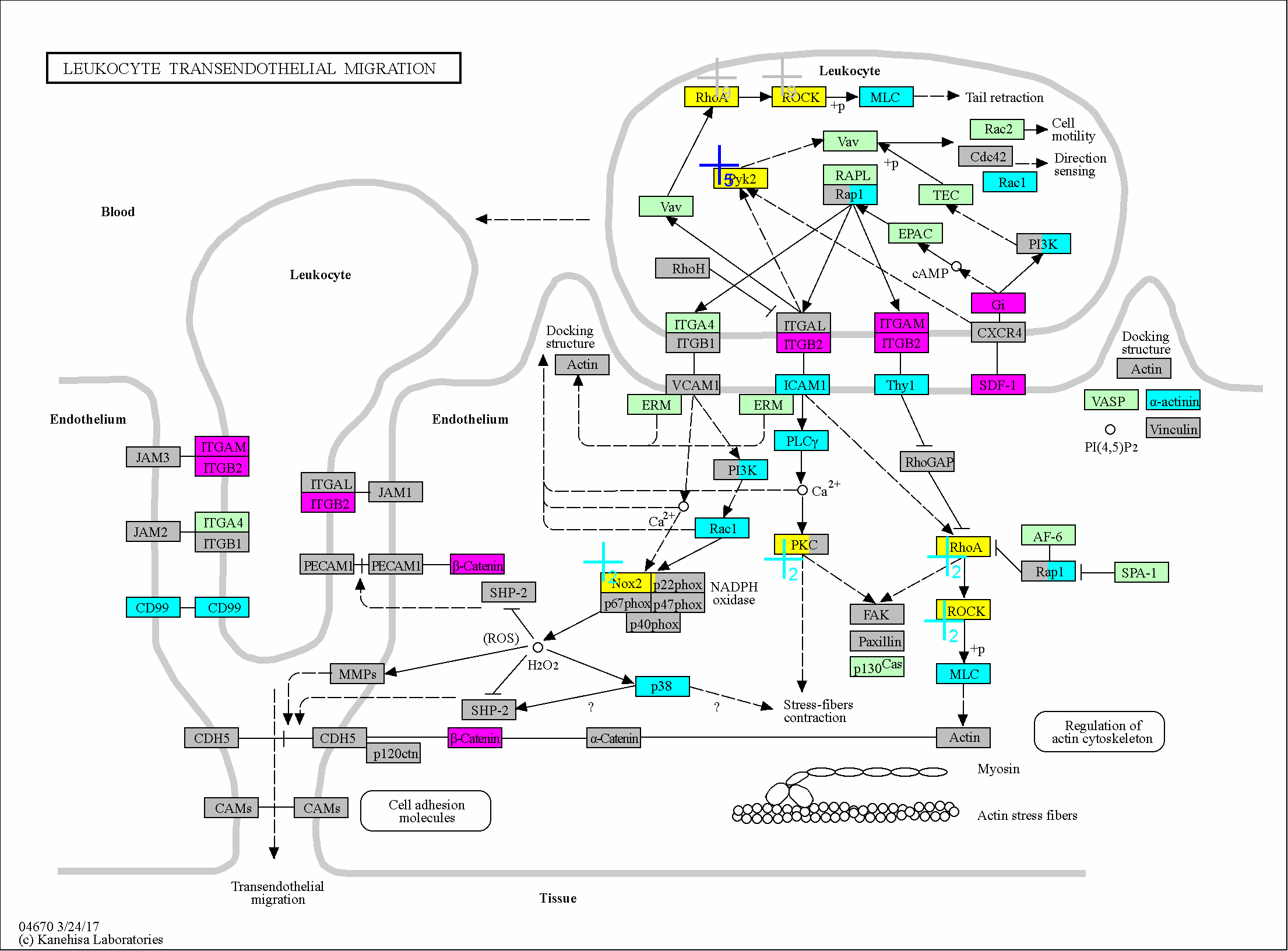

### 4720

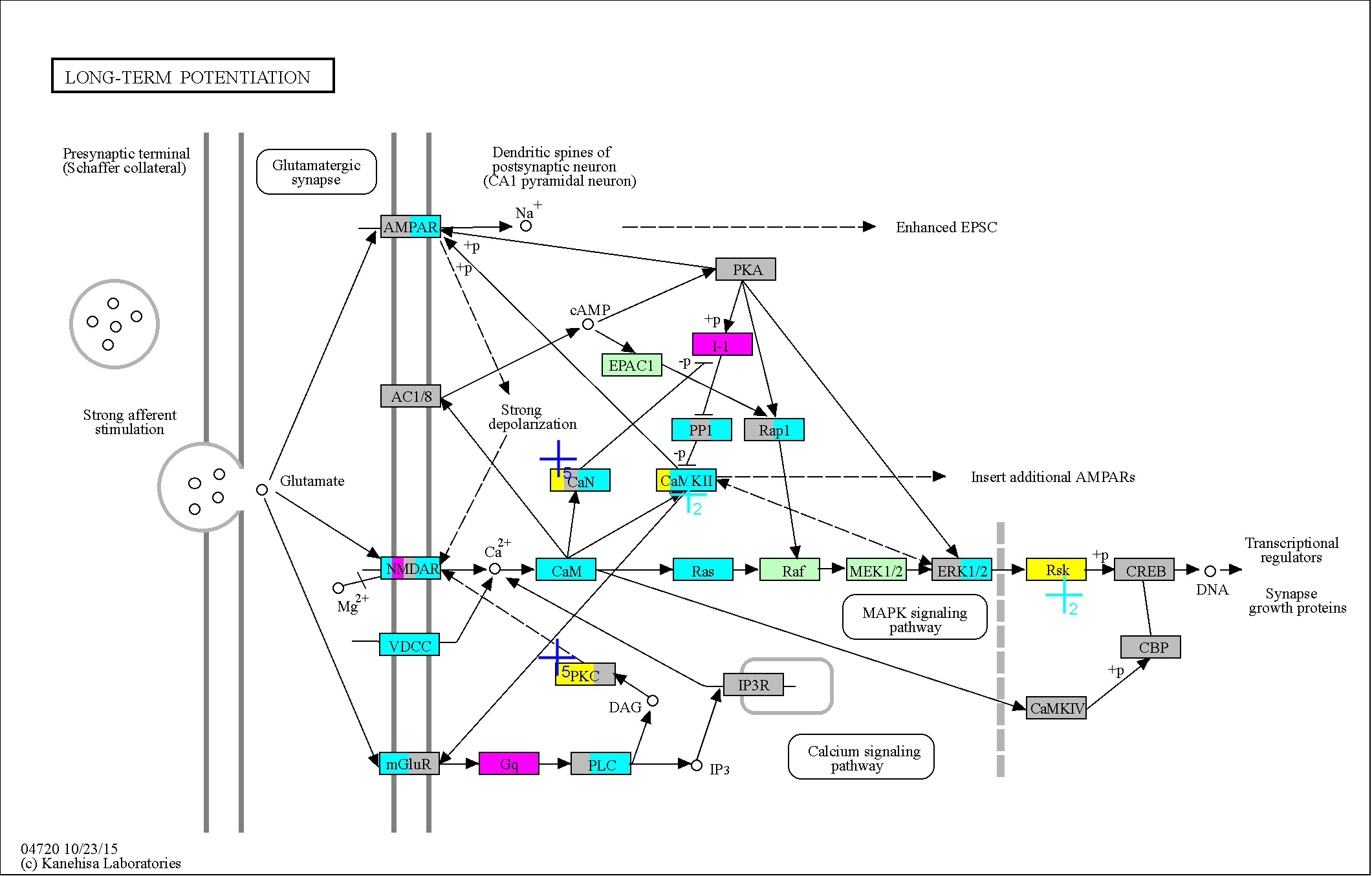

### 4722

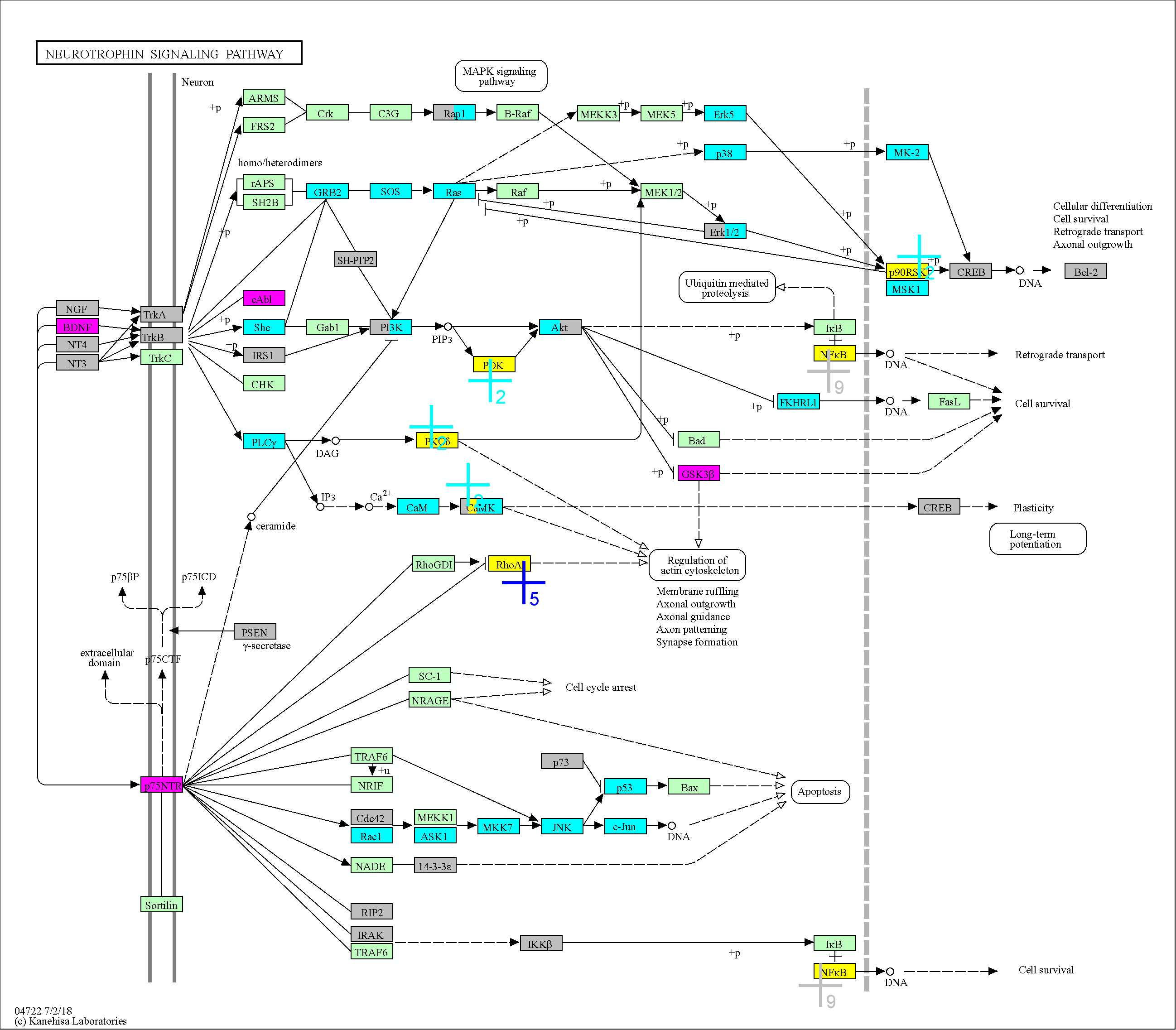

### 4723

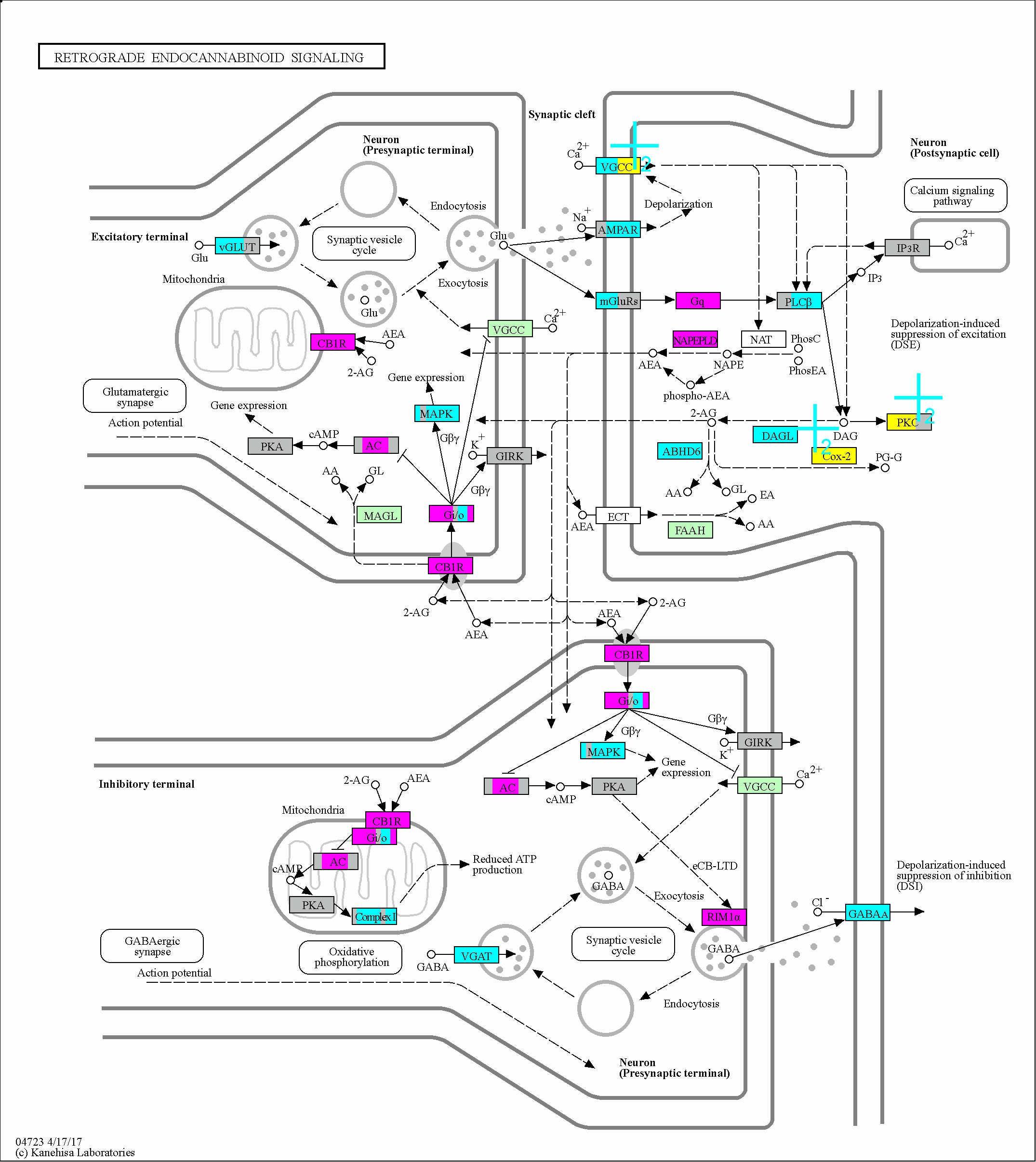

### 4724

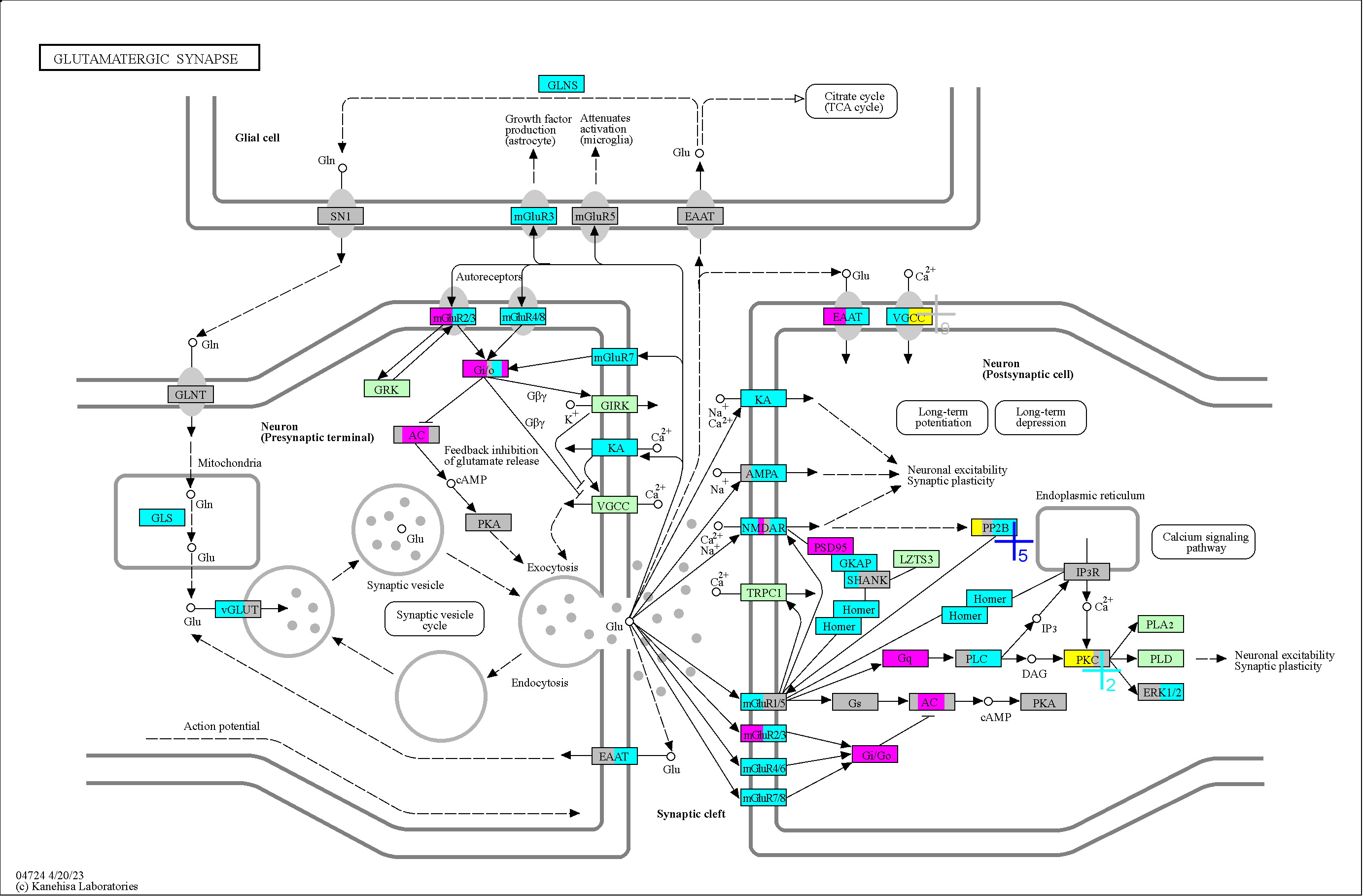

### 4725

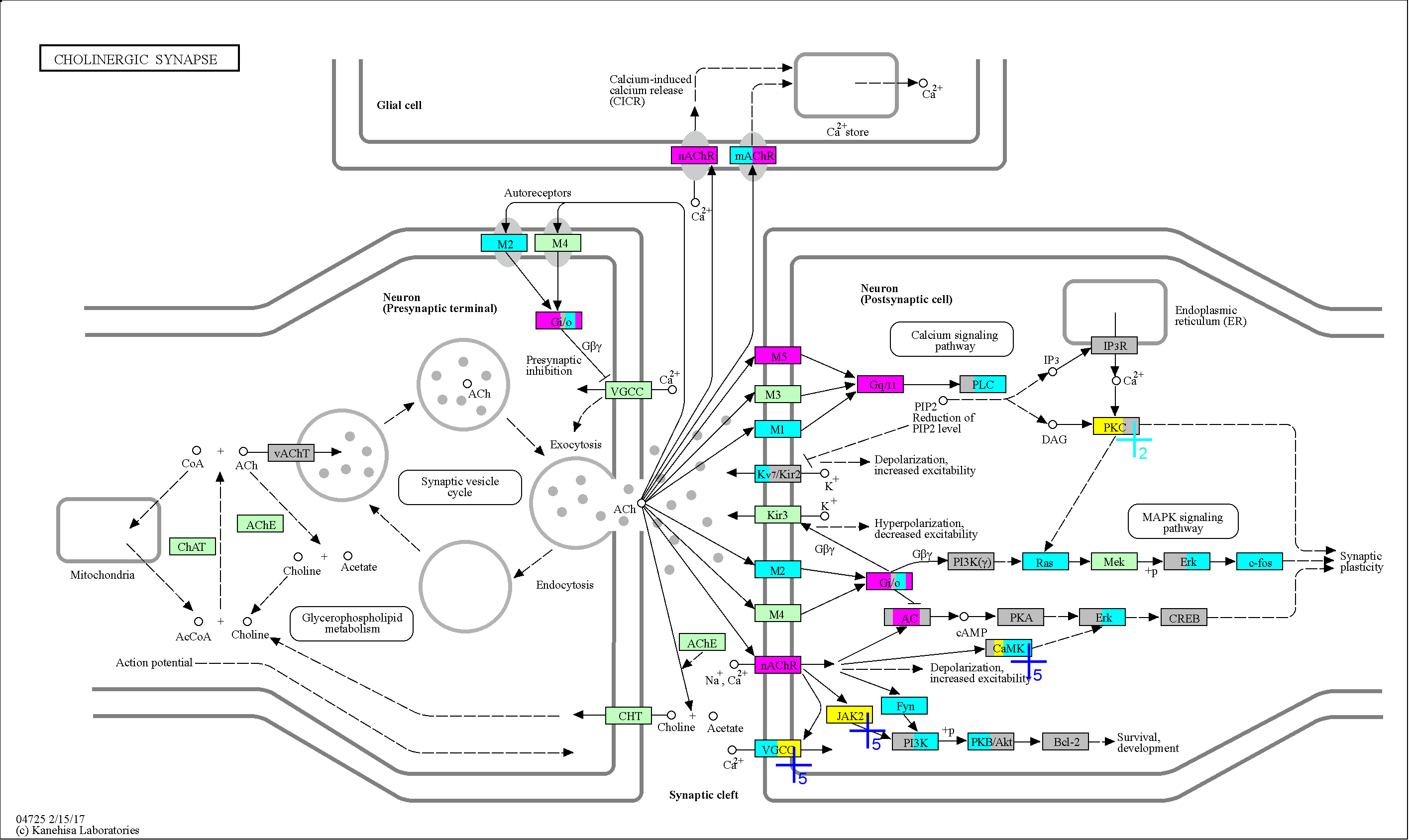

### 4726

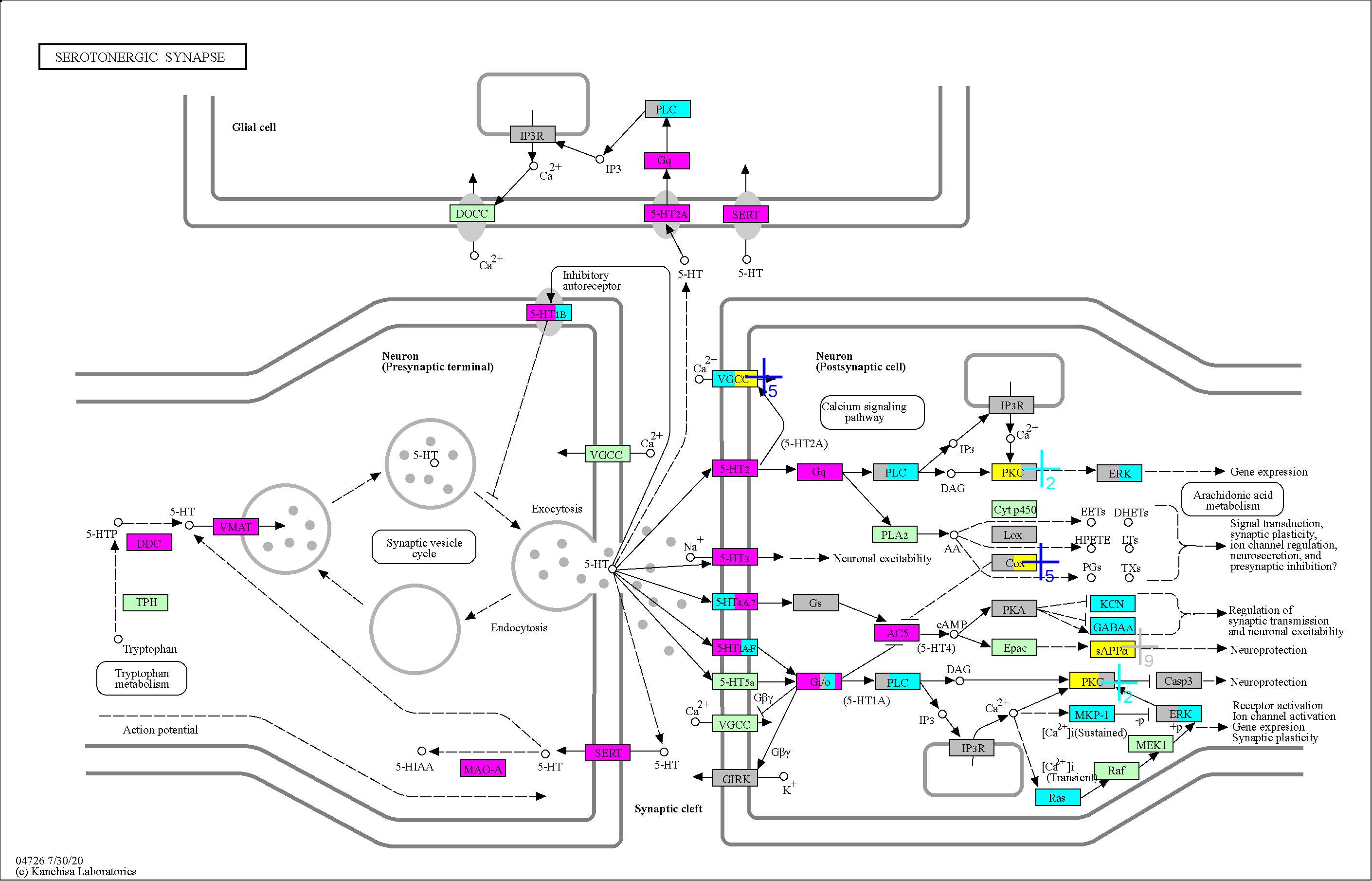

### 4727

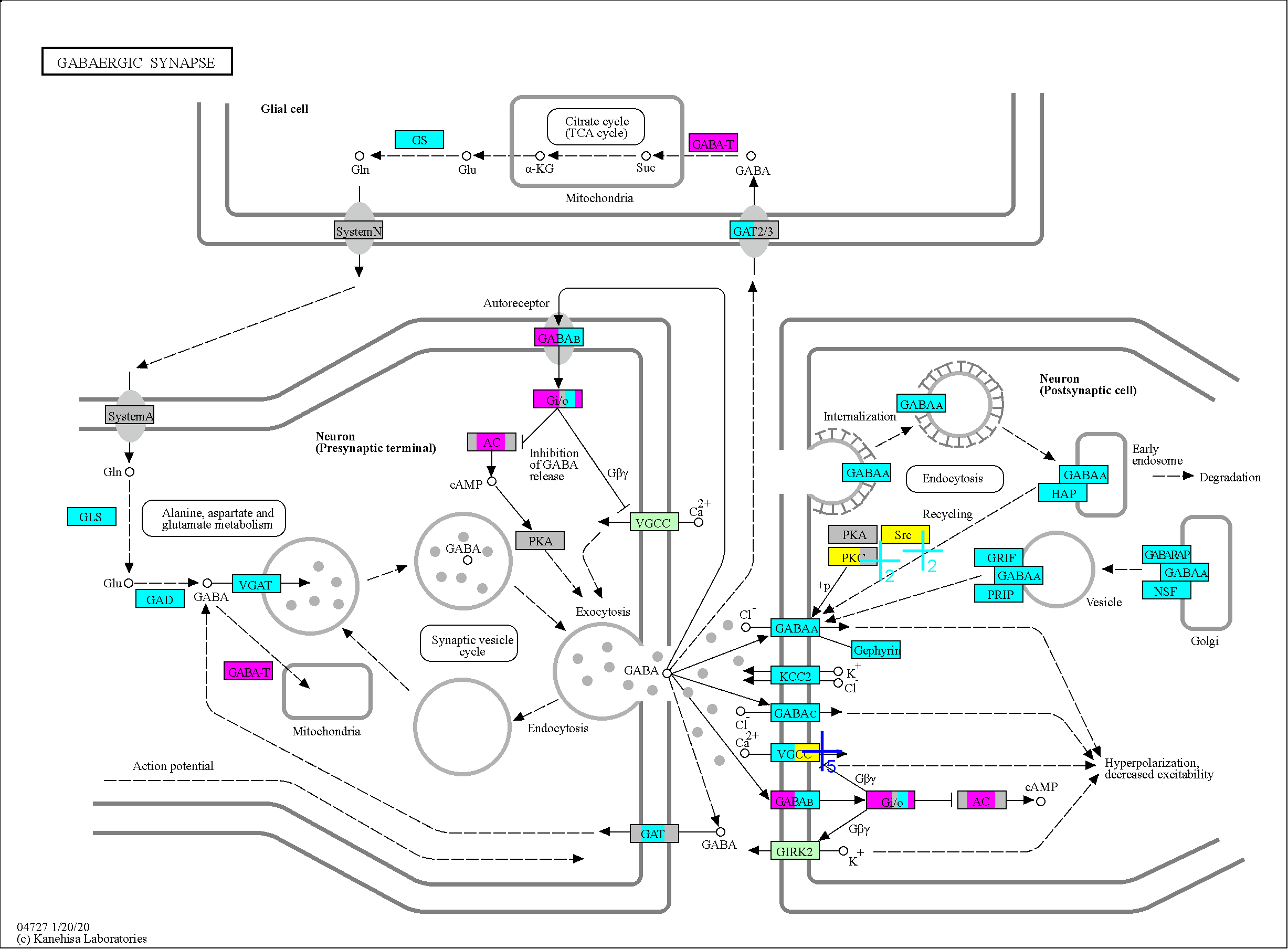

### 4728

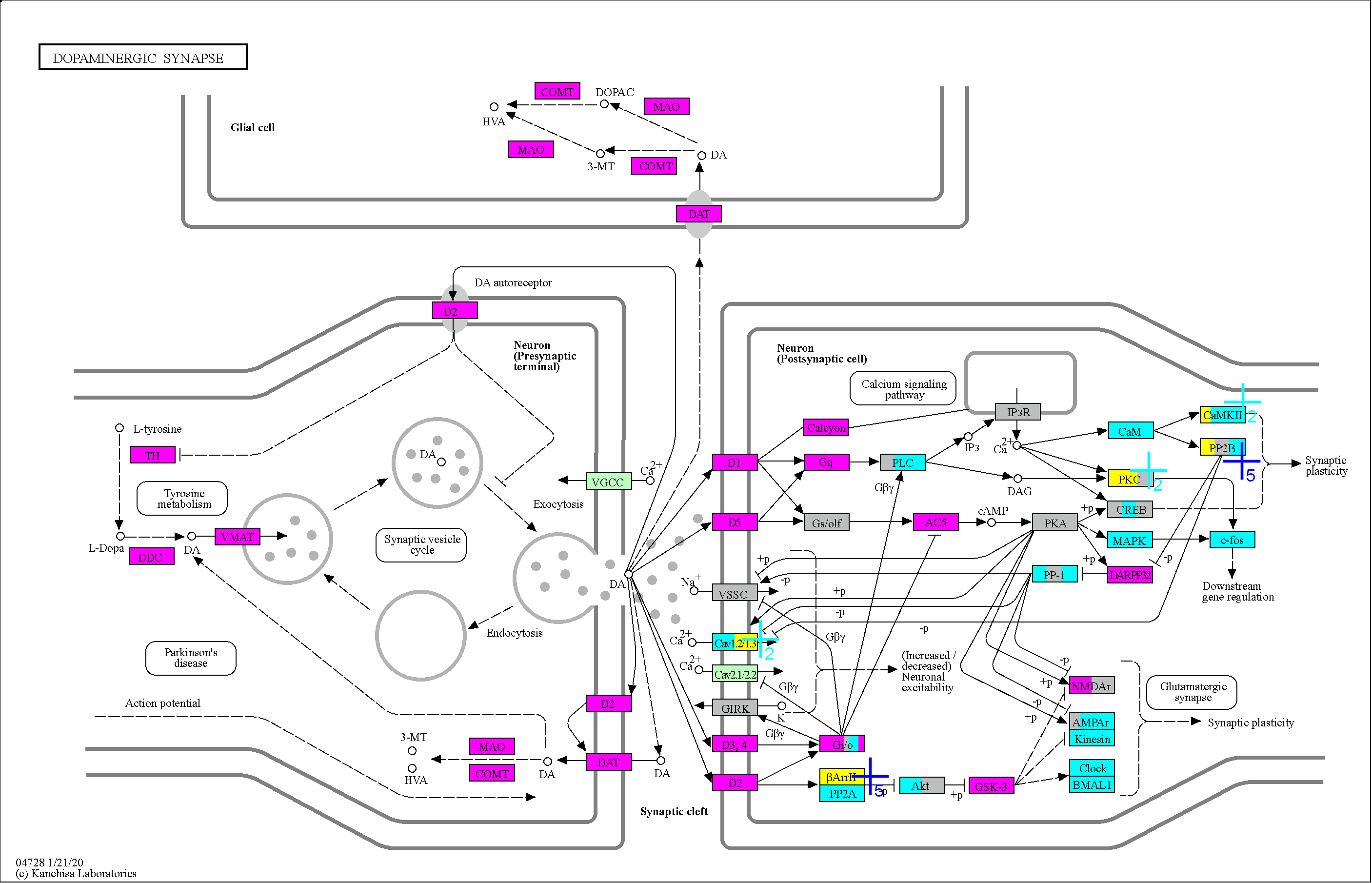

### 4750

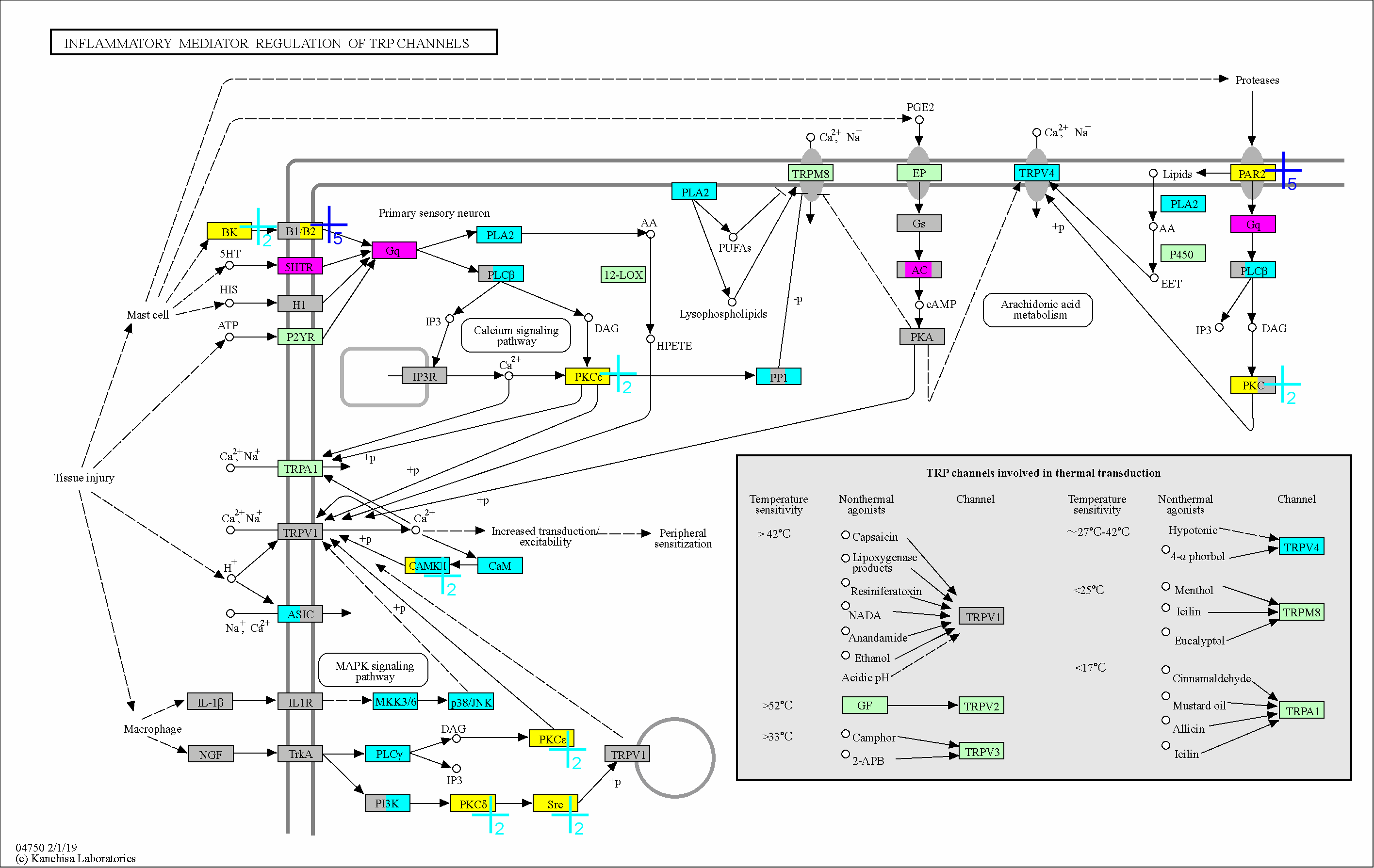

### 4810

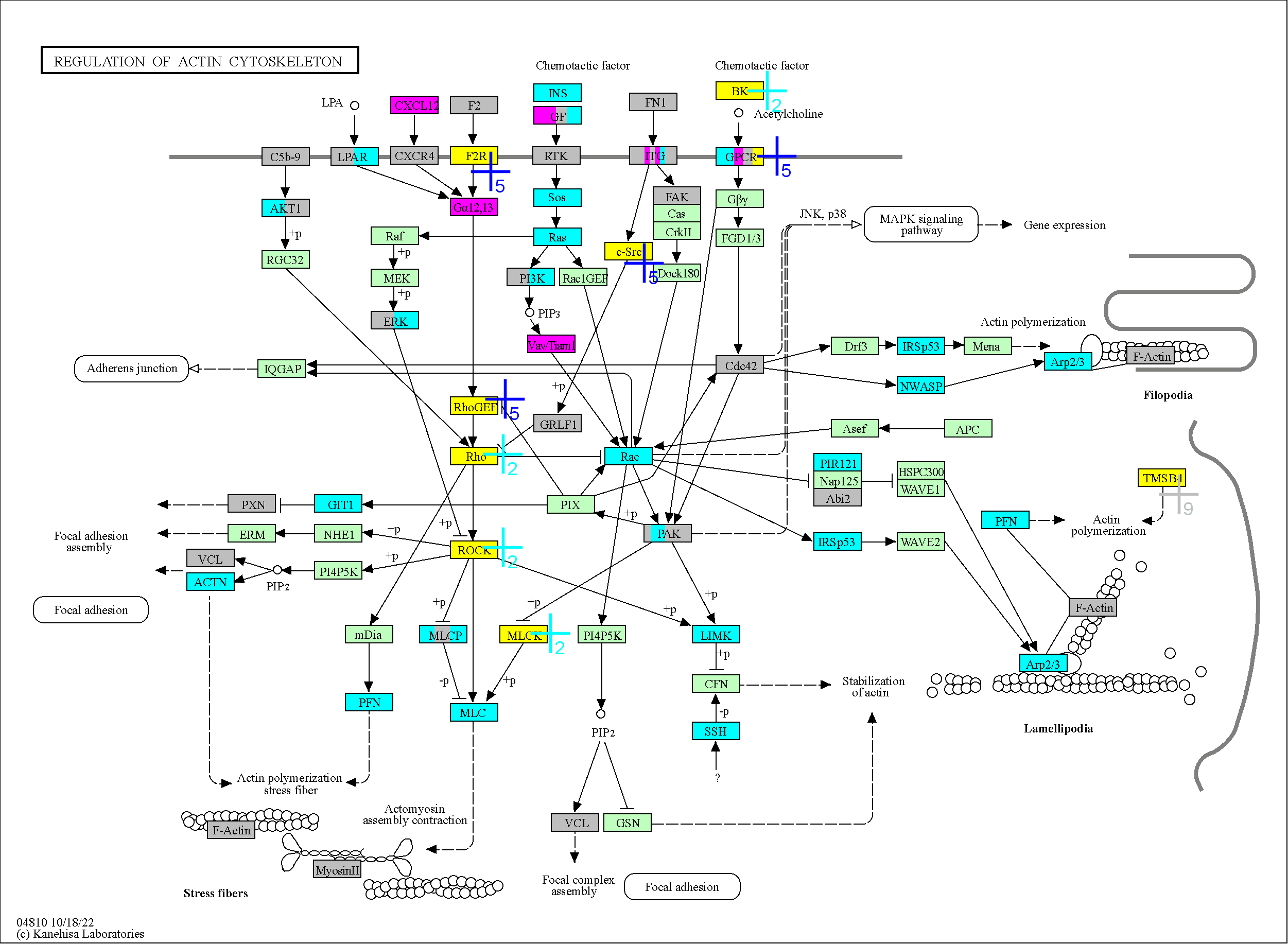

### 4911

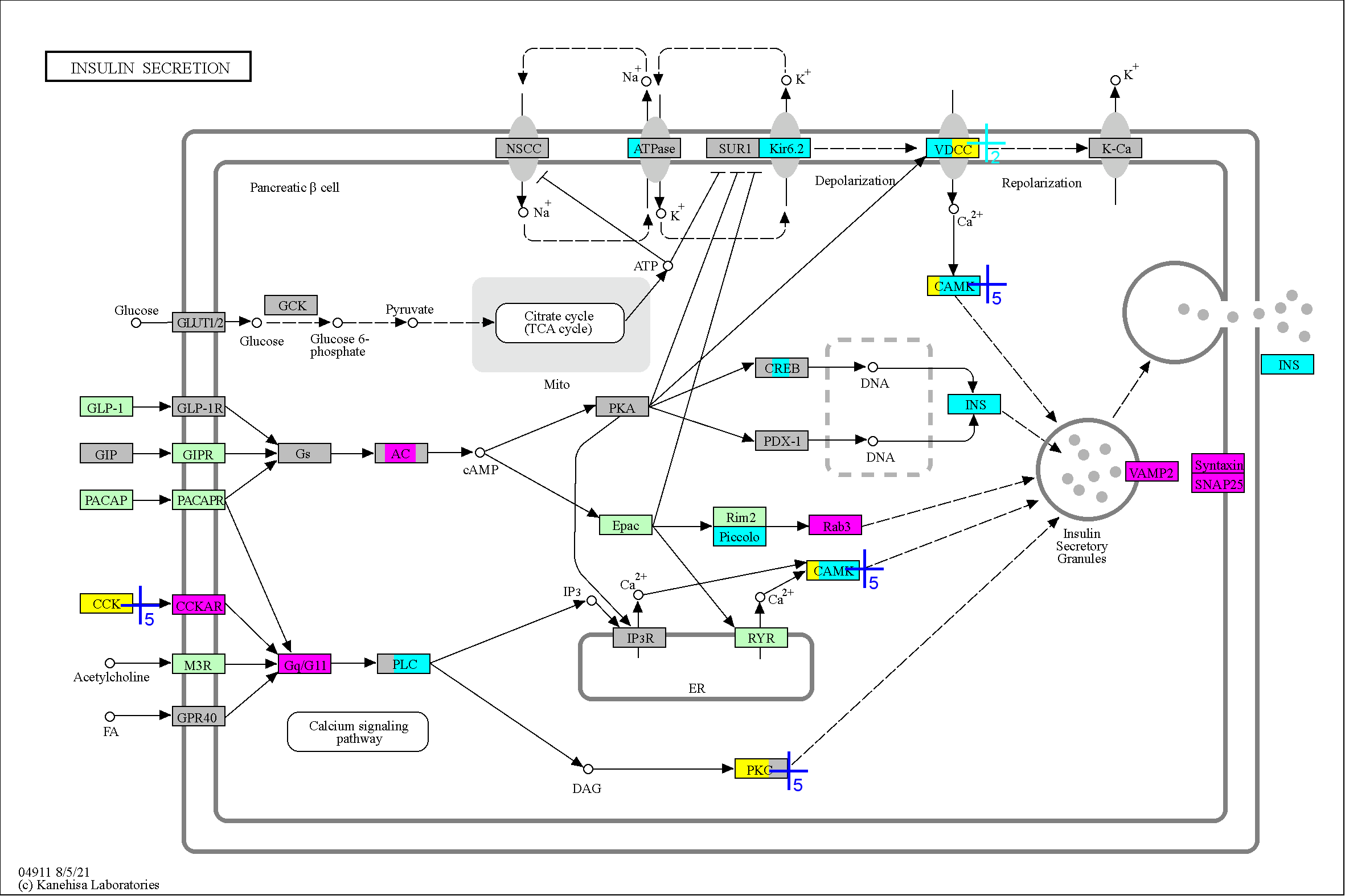

### 4912

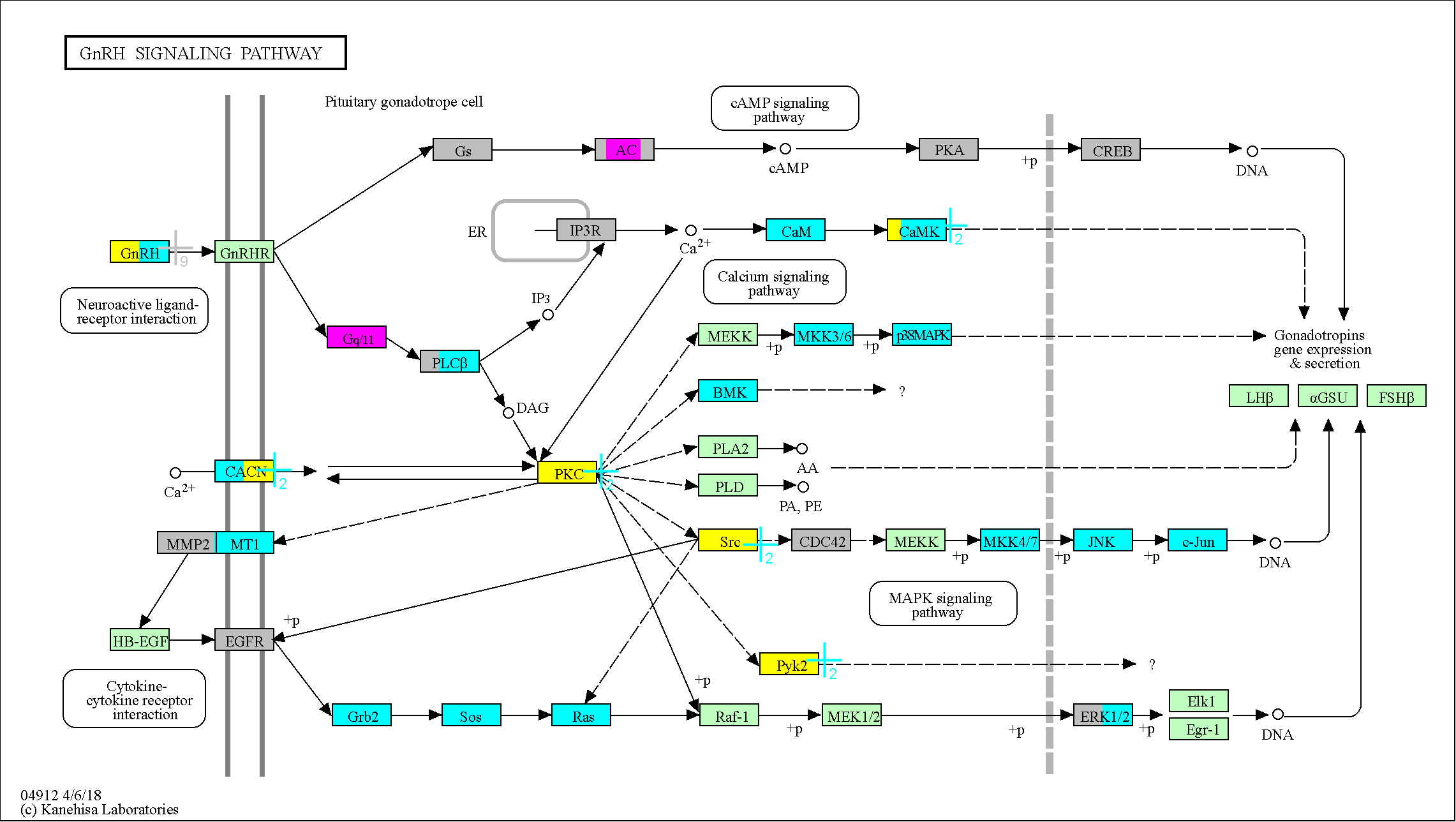

### 4913

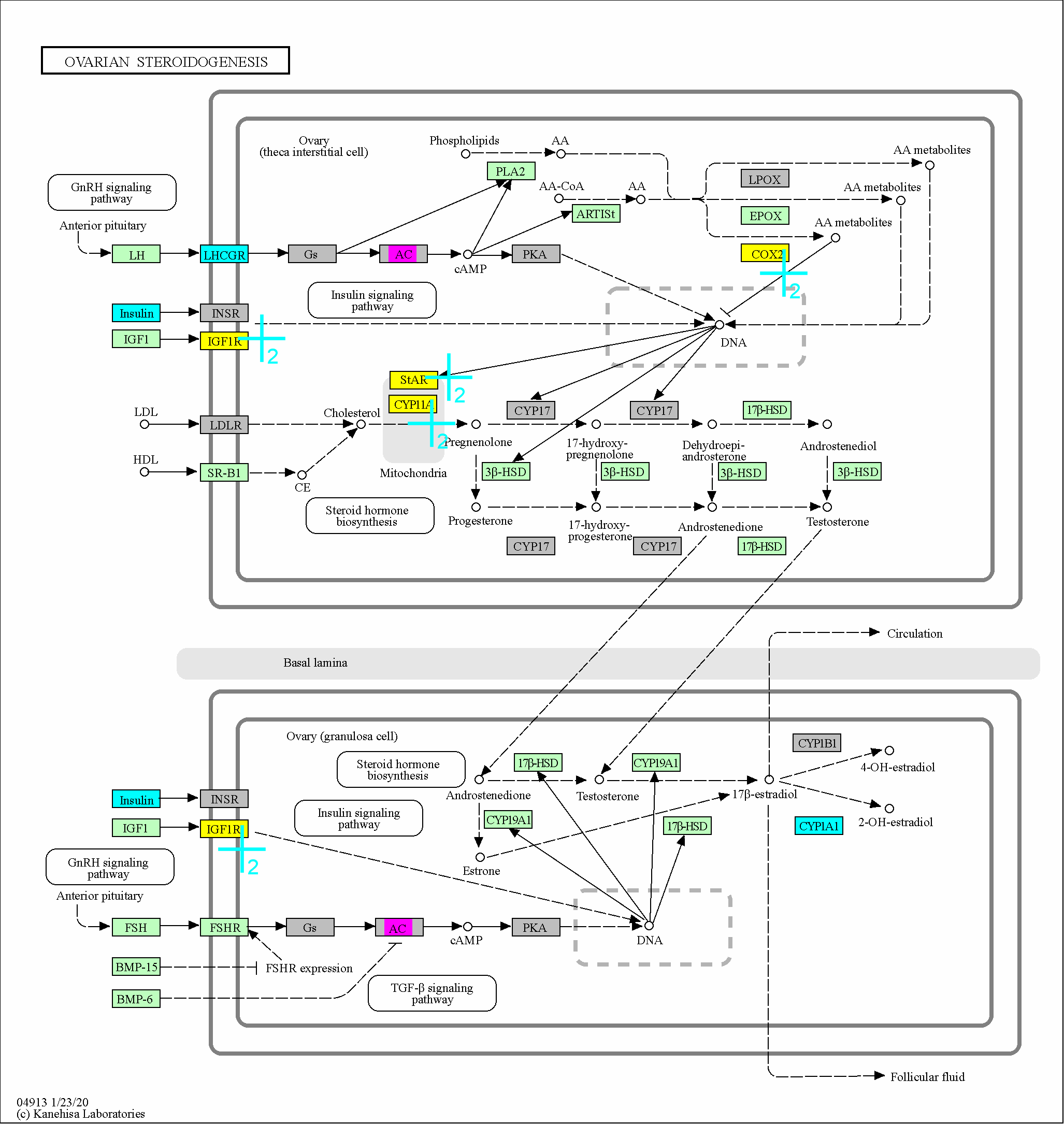

### 4916

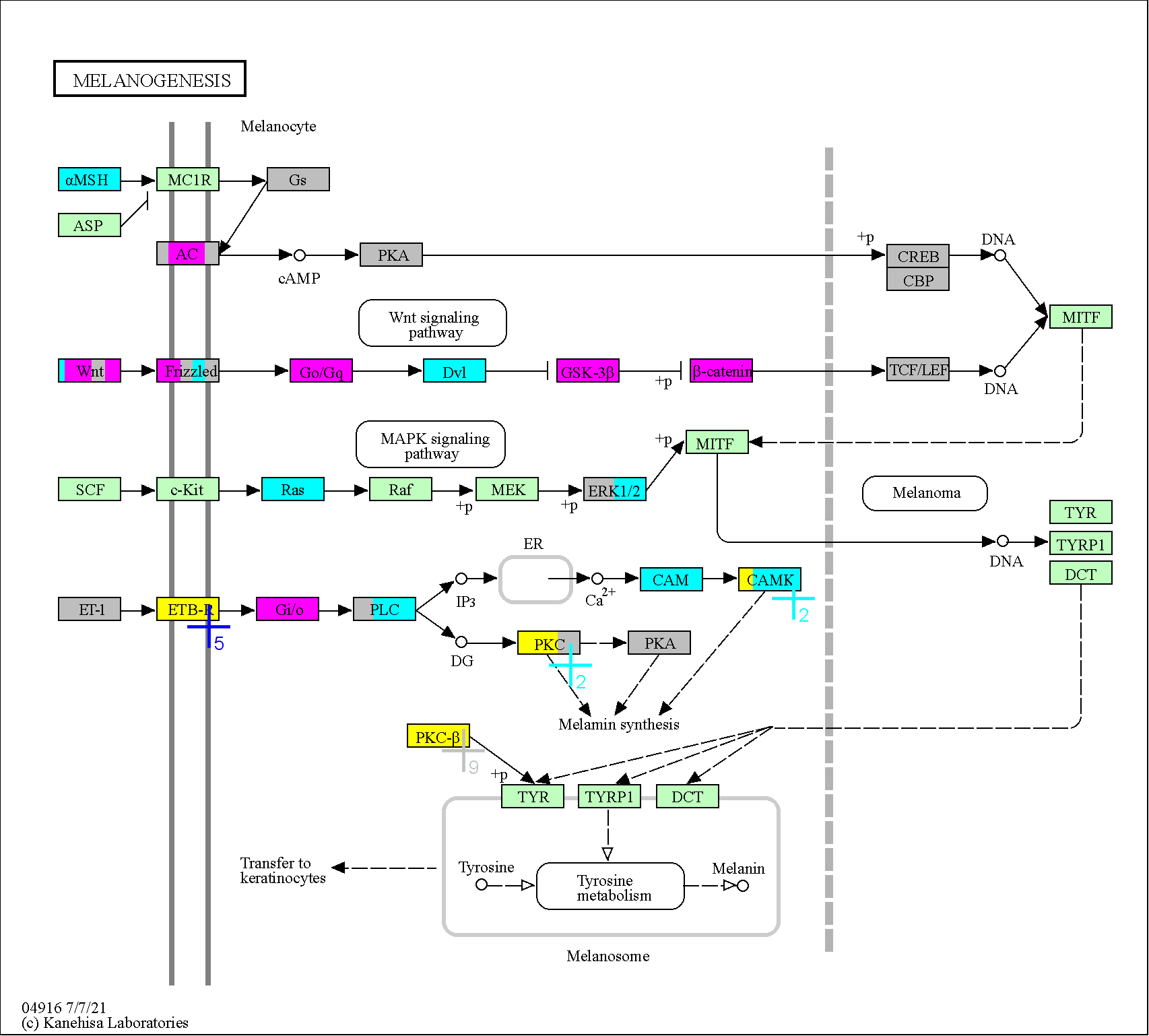

### 4918

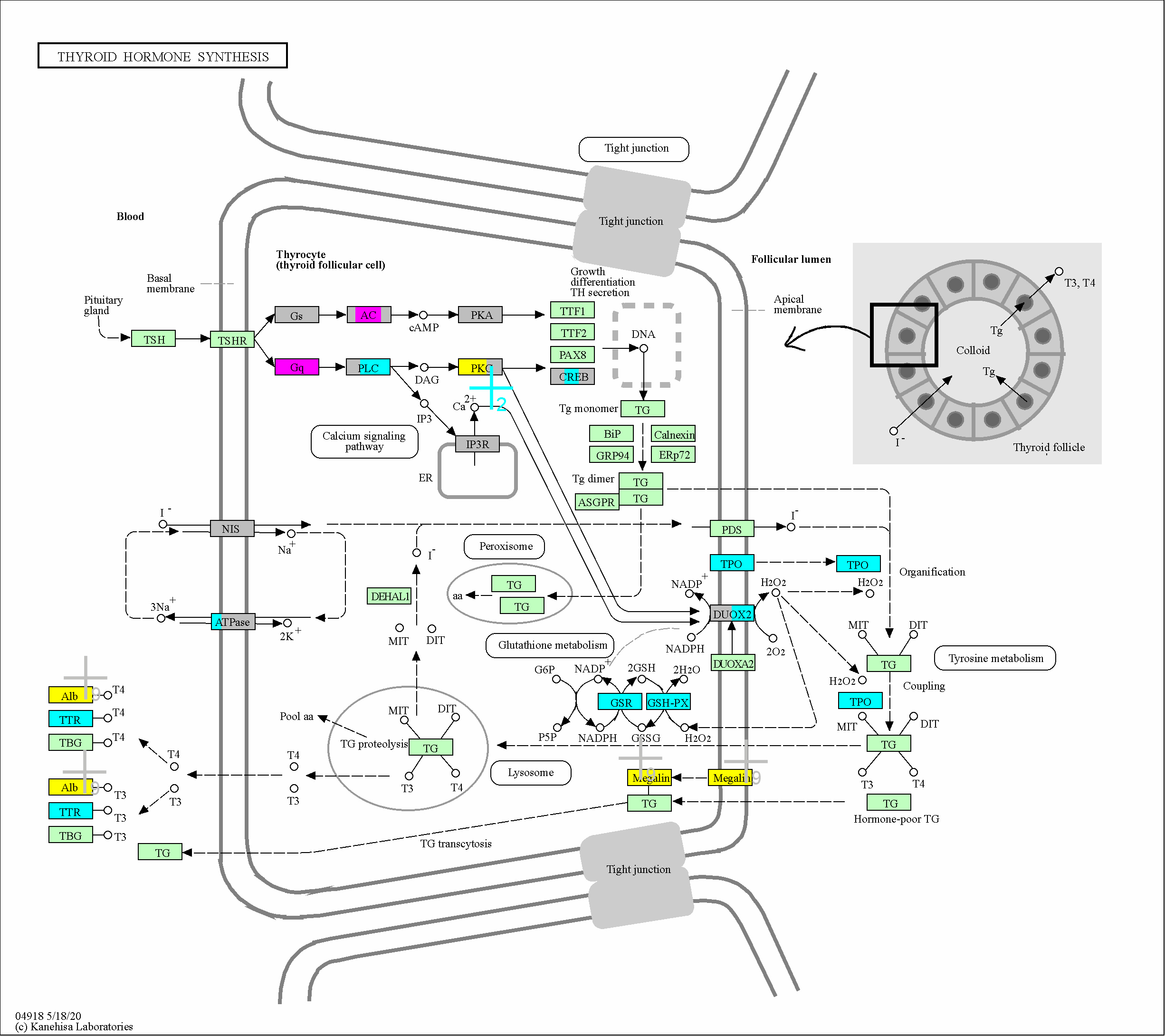

### 4919

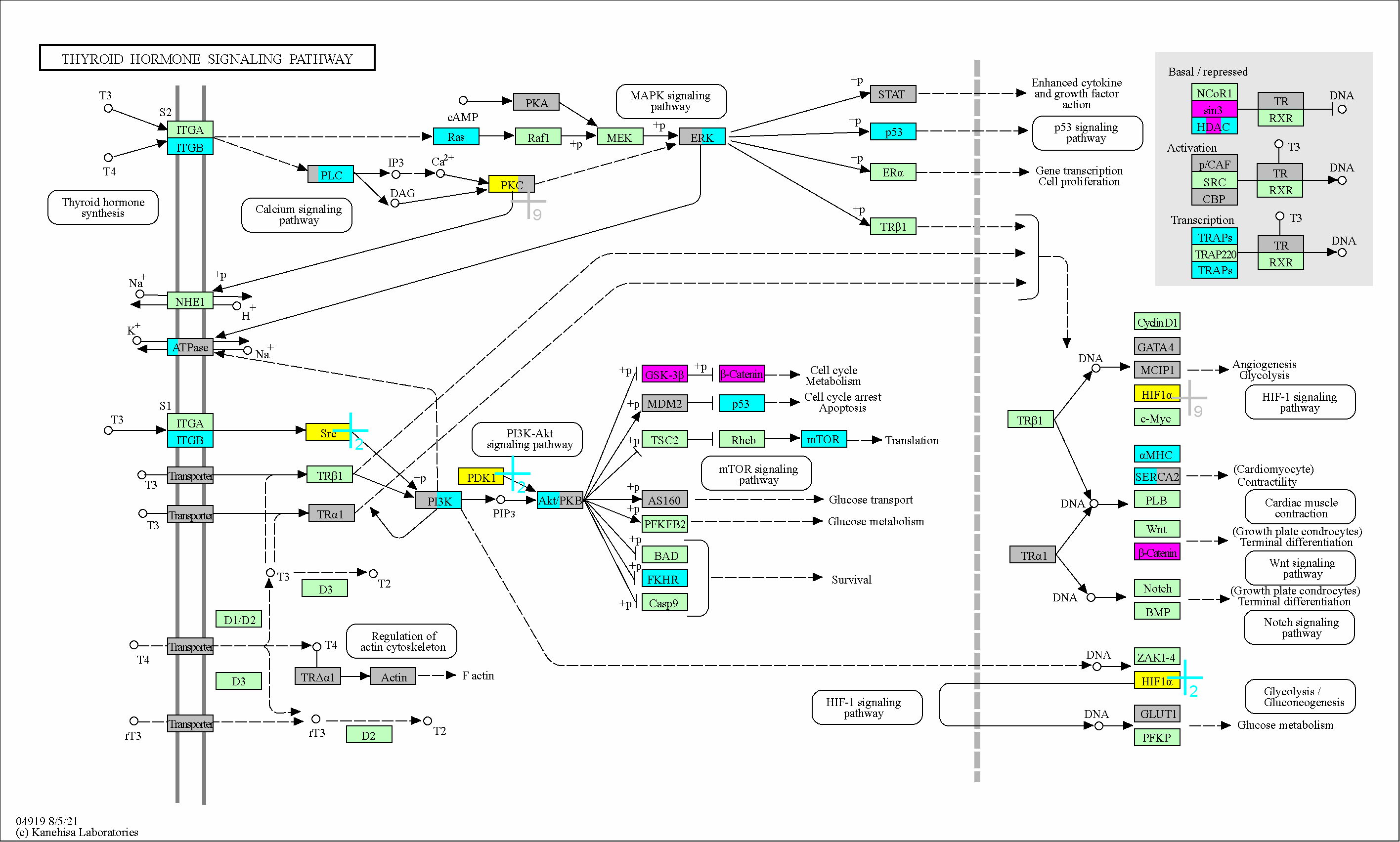

### 4921

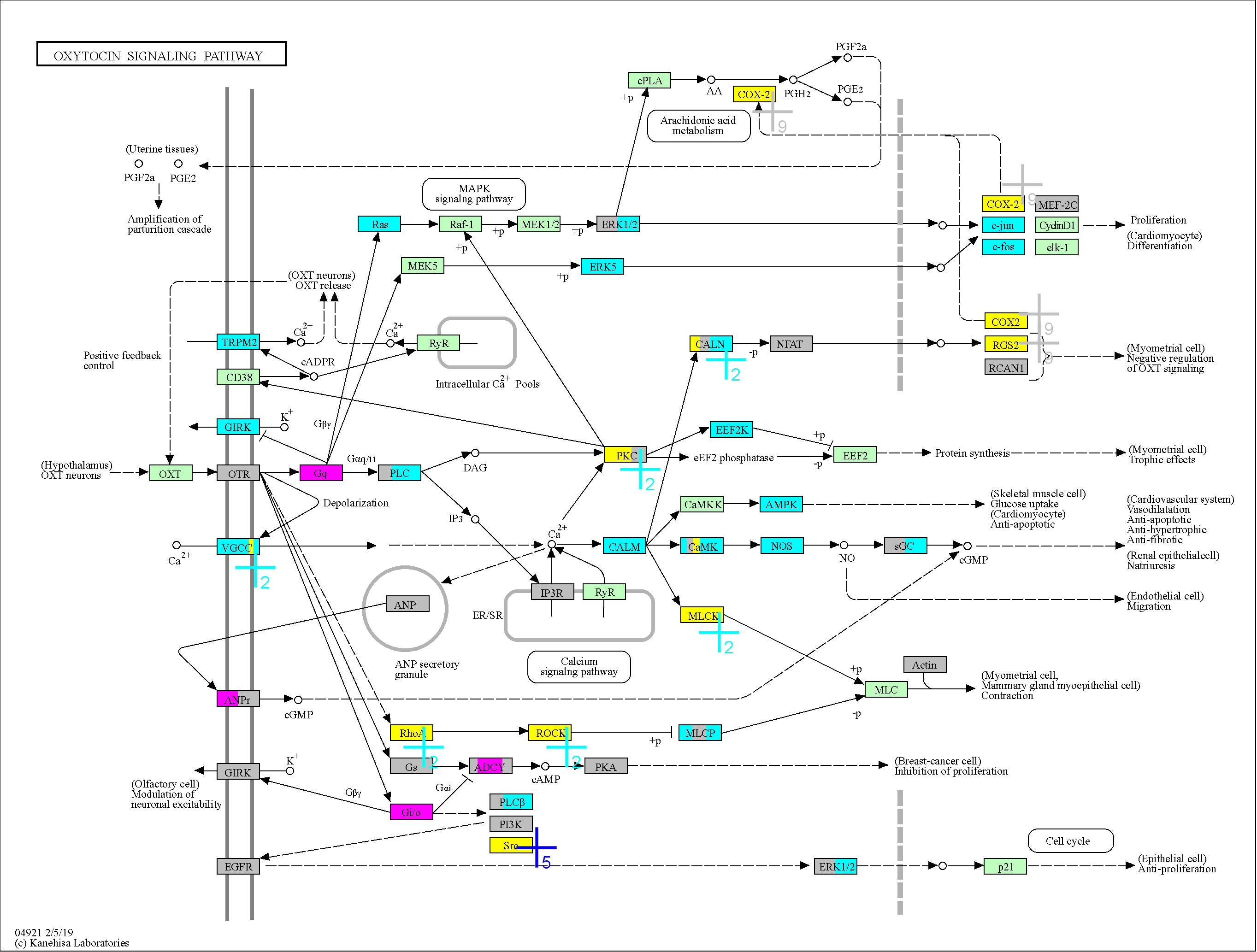

### 4924

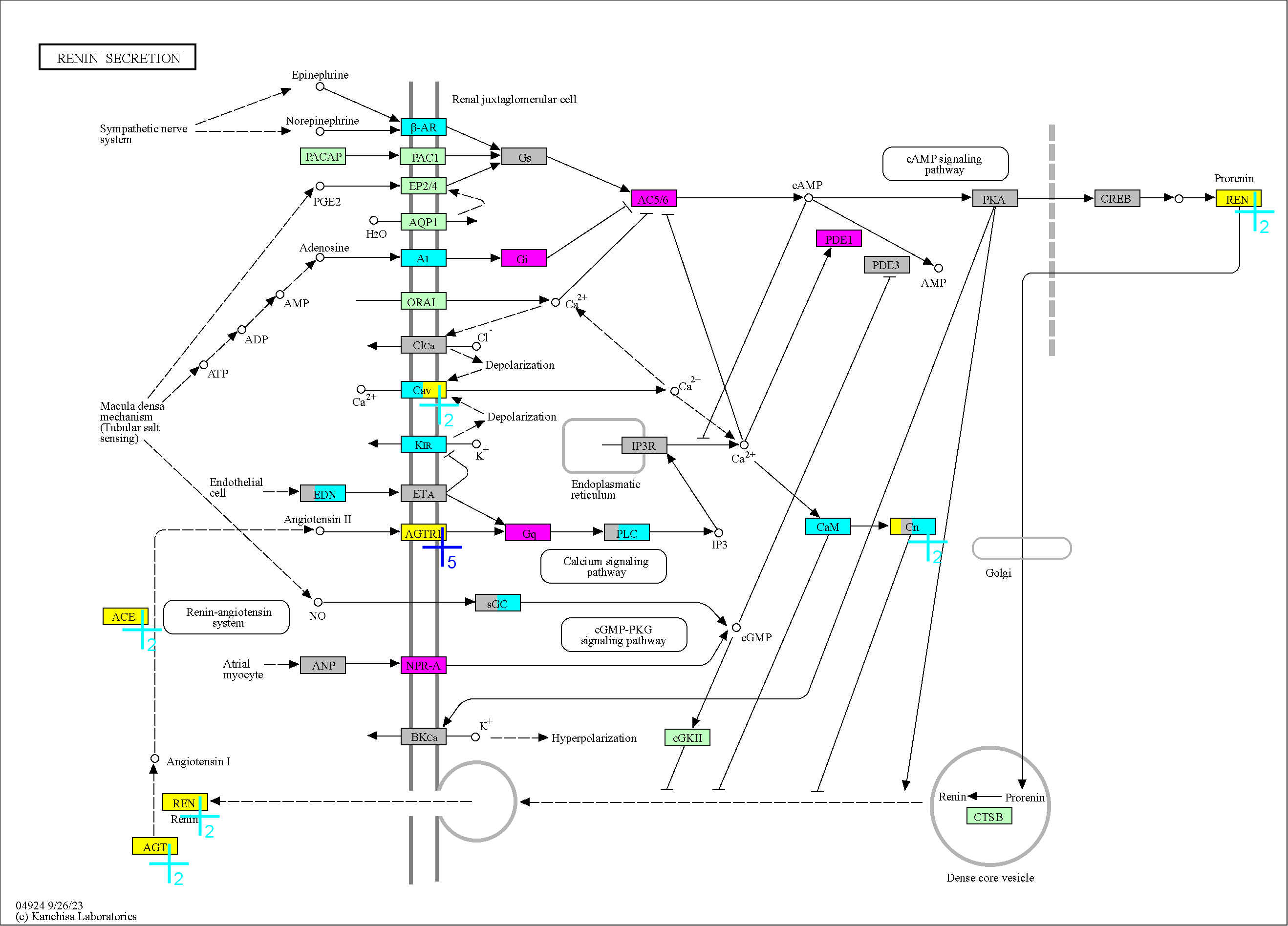

### 4925

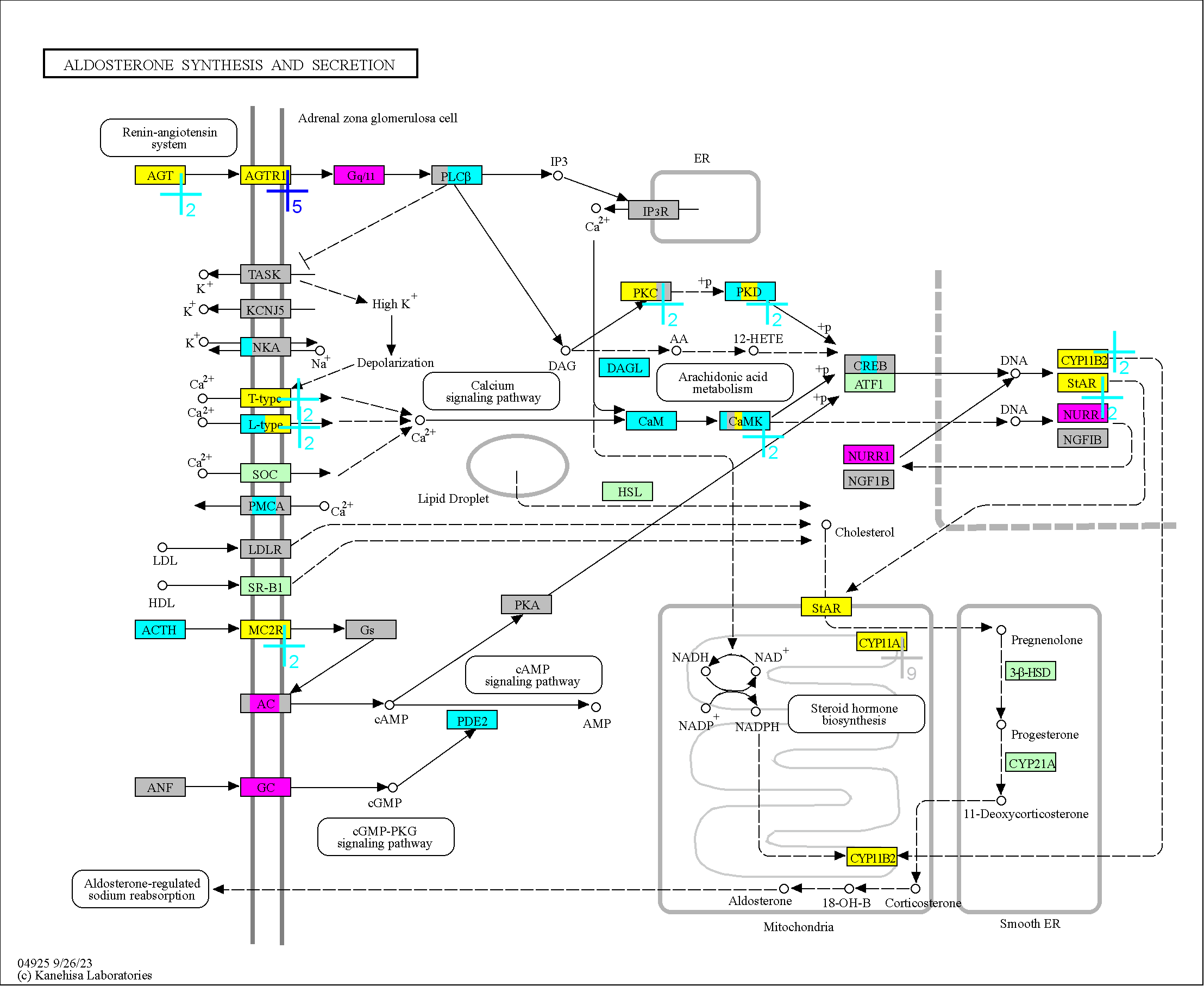

### 4926

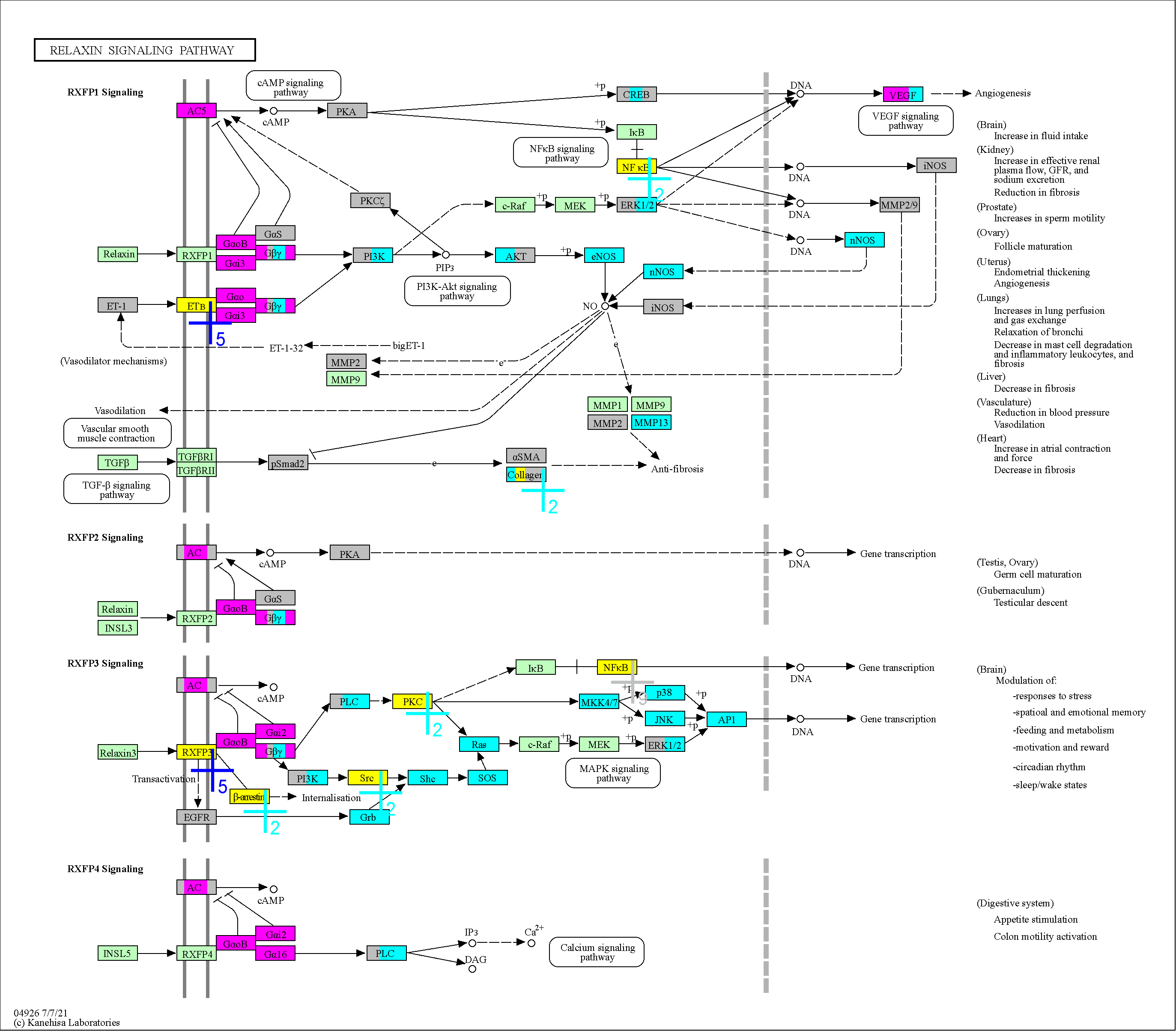

### 4927

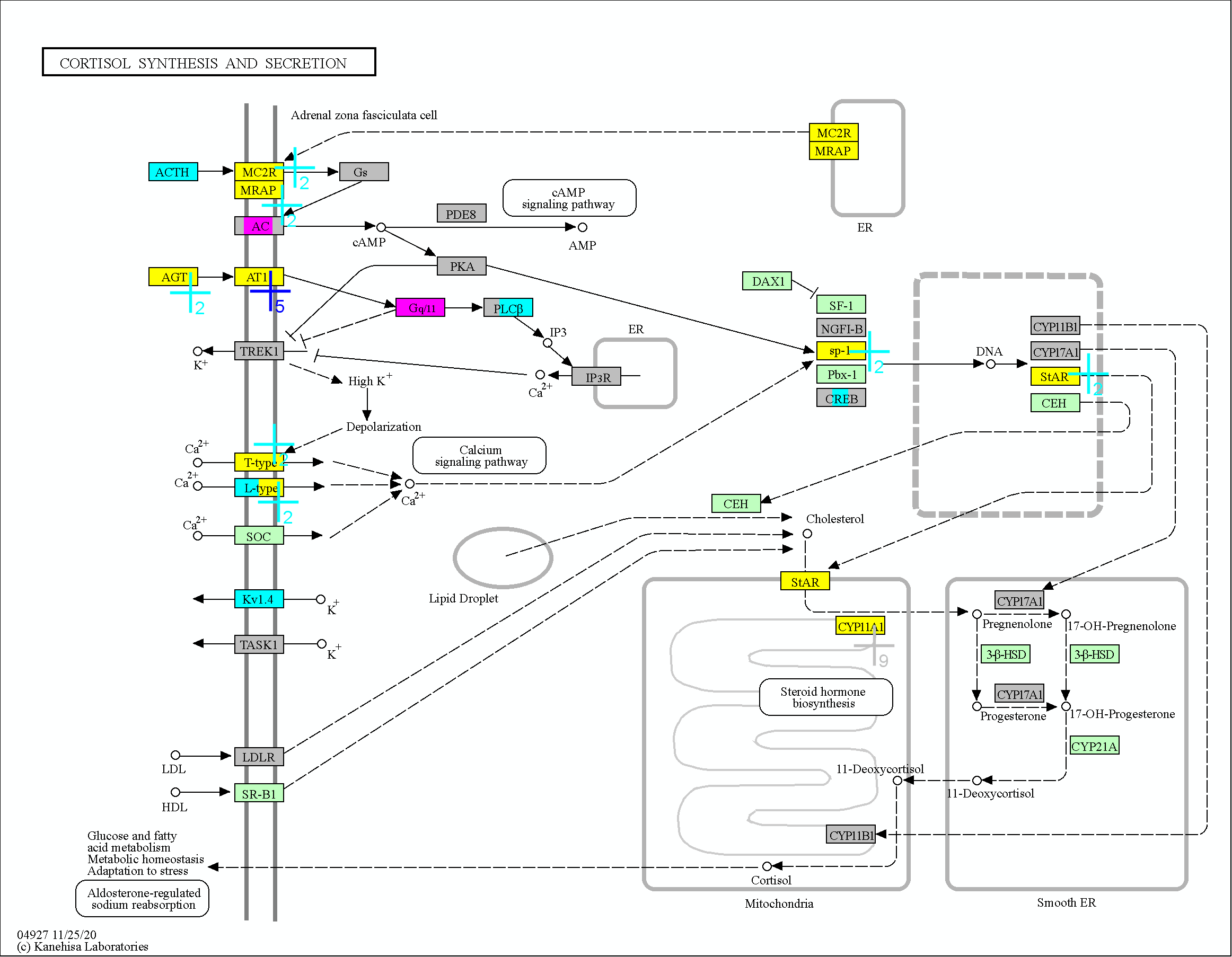

### 4928

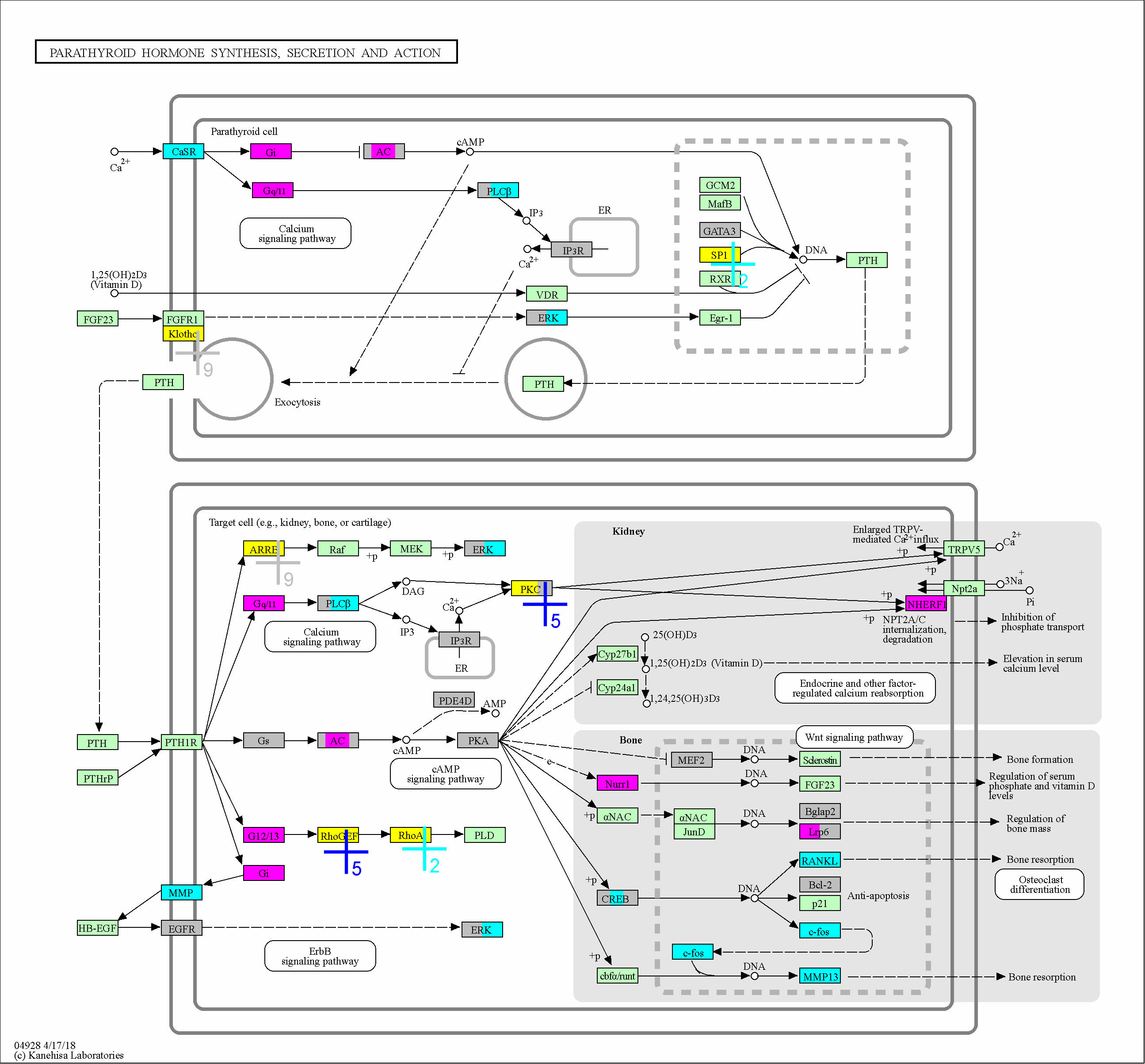

### 4929

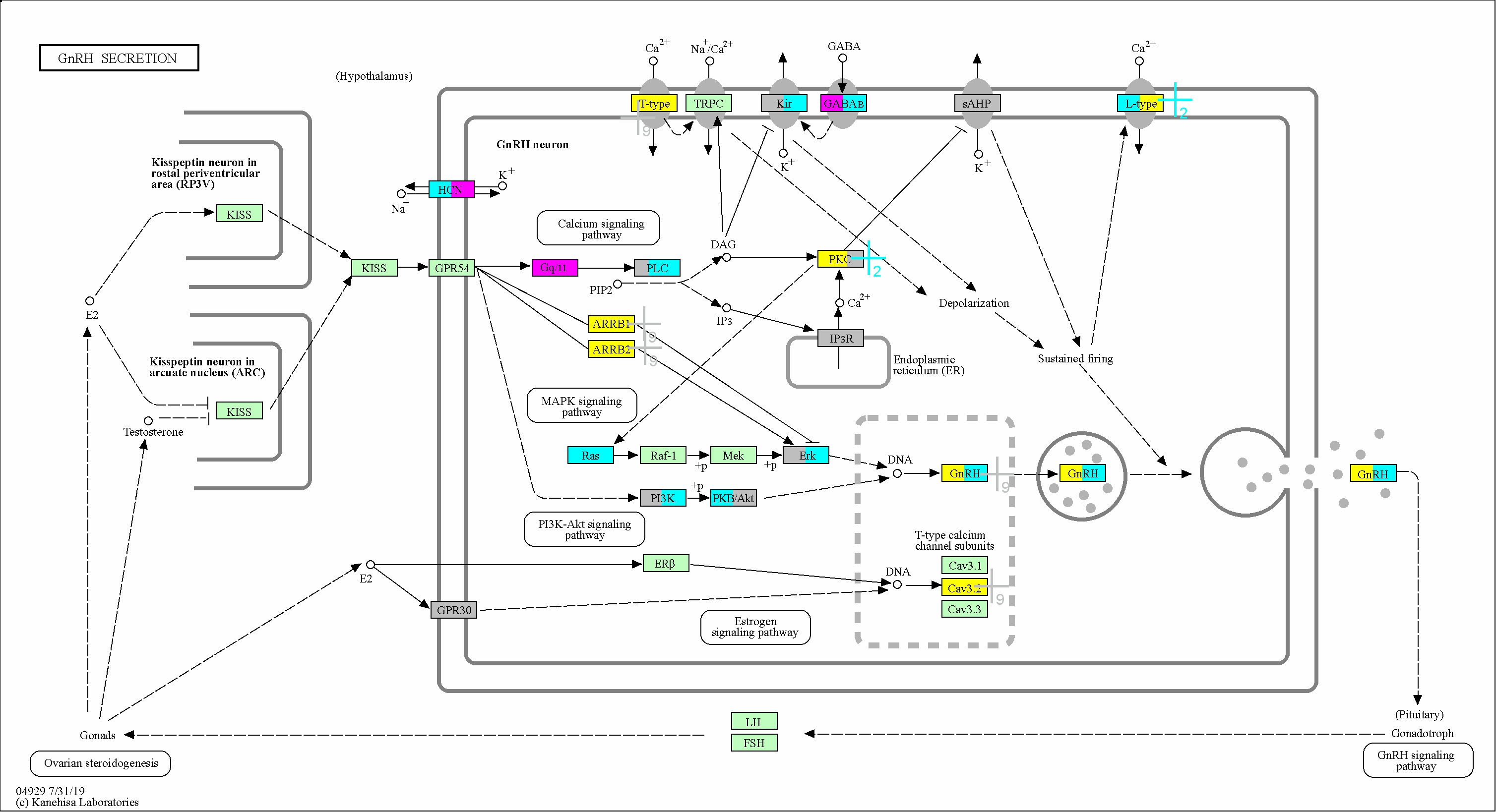

### 4935

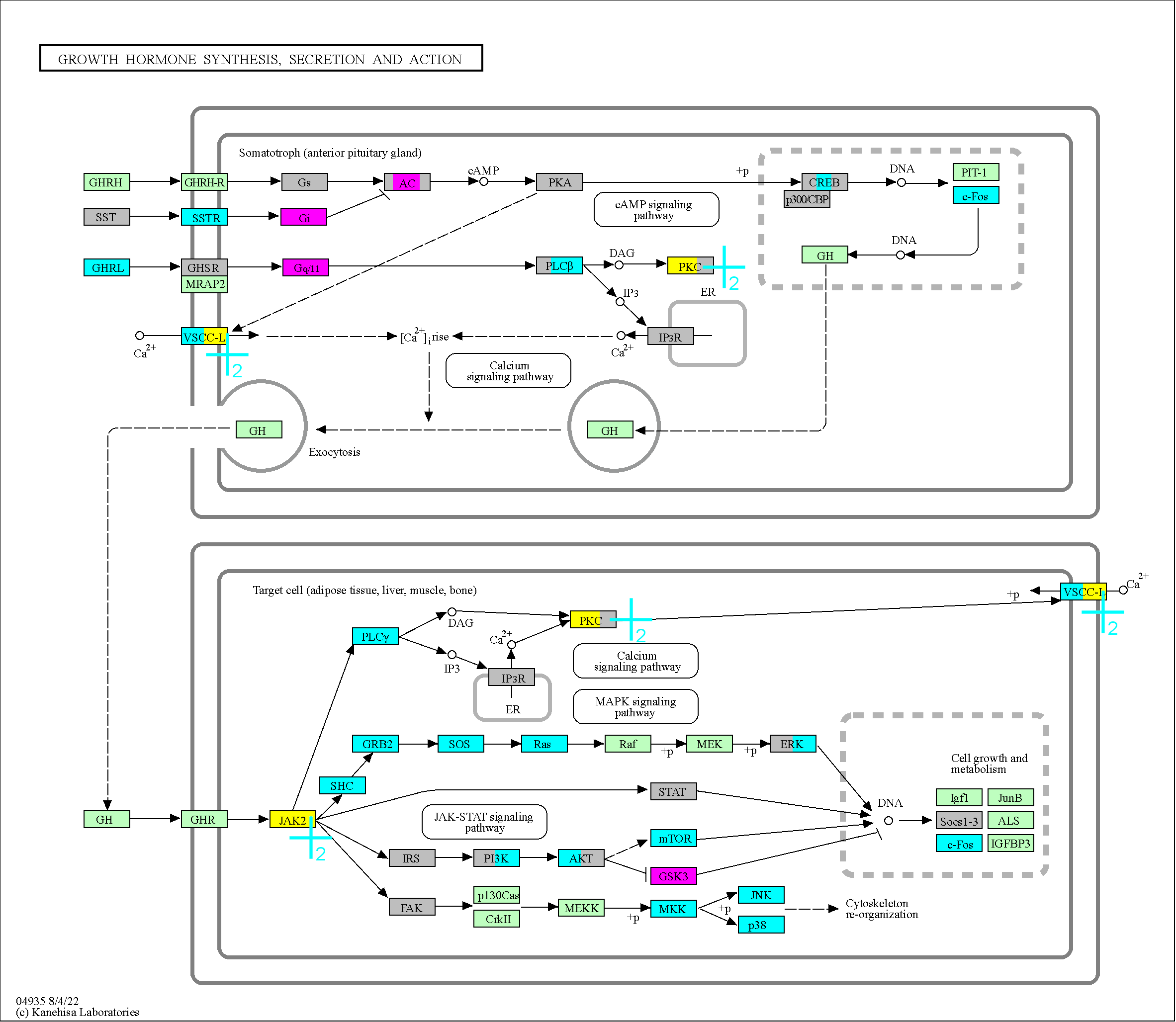
